# AI-driven discovery of host thioredoxin as a CRISPR enhancer of phage-encoded miniature Cas12 hacker nuclease

**DOI:** 10.1101/2025.01.20.633832

**Authors:** Zhipeng Wang, Yujue Wang, Hui Gao, Jiani Dai, Na Tang, Yannan Wang, Quanjiang Ji

**Affiliations:** School of Physical Science and Technology & State Key Laboratory of Advanced Medical Materials and Devices, ShanghaiTech University, Shanghai, 201210, China; Gene Editing Center, School of Life Science and Technology, ShanghaiTech University, Shanghai, 201210, China; Shanghai Clinical Research and Trial Center, Shanghai, 201210, China

## Abstract

The evolutionary arms race between bacteriophages and their bacterial hosts has driven the evolution of sophisticated adaptive immune systems, such as CRISPR-Cas, as a crucial defense mechanism. While bacteriophages have developed various anti-CRISPR strategies to counteract these immune systems, the role of bacterial host factors in enhancing CRISPR-Cas functions has been relatively unexplored. In this study, we employ an artificial intelligence (AI)-driven approach to systematically analyze potential interactions between *Escherichia coli* (*E. coli*) proteins and fifteen previously uncharacterized Cas12 proteins, generating 65,715 predicted binary complex structures. Our findings reveal a previously unknown dimension of CRISPR immunity, demonstrating that the host’s ubiquitous redox enzyme, thioredoxin (TrxA), significantly enhances the DNA cleavage efficiency of a phage-encoded, miniature Cas12 nuclease (termed ‘Cas12 hacker’). This synergistic relationship represents a strategic inversion, where a bacteriophage hijacks a host protein to reinforce its own genome degradation machinery, possibly targeting rival nucleic acids. Through comprehensive biochemical characterizations, structural analyses of the Cas12 hacker-TrxA-sgRNA-dsDNA quaternary complex, and *in vivo* bacterial defense assays, we uncover an intricate association in which thioredoxin binds to and activates the Cas12 hacker nuclease, intensifying its DNA cleavage capacity and bolstering CRISPR immunity. Our findings expand the understanding of the molecular interactions underlying host-phage conflicts and highlight the potential for harnessing endogenous host factors to enhance the capabilities of CRISPR-based genetic engineering tools.

## Introduction

The antagonistic coevolution of bacteria and Mobile Genetic Elements (MGEs) is characterized by a dynamic series of defense-counterdefense strategies. Among these, the CRISPR-Cas system exemplifies a critical bacterial defense mechanism, enabling the host to counteract exogenous MGEs^1–5^. Conversely, MGEs, notably bacteriophages, have evolved sophisticated countermeasures to circumvent CRISPR-Cas-mediated defenses. These include the production of Anti- CRISPR (Acr) proteins and RNAs that inhibit Cas effector function or disrupt CRISPR complex assembly, representing a significant challenge to bacterial immunity^6–12^. In response, bacteria have developed advanced counter-strategies, such as type I-F CRISPR systems that trigger abortive infection-like mechanisms to eliminate Acr-containing elements^13^. Moreover, some type II-C CRISPR systems utilize pro-CRISPR proteins to amplify their MGE-targeting efficacy^14^.

The Cas12 family encapsulates a prime example of the coevolutionary complexity inherent in these host-phage interactions, demonstrating remarkable versatility across its various subclasses^15,16^. Notably diverse in structure, the key domains (wedge (WED), recognition (REC), and RuvC) span a wide range of sizes^17^, accommodating proteins from 400 to 1,500 amino acids, which underlines the adaptive potential of these systems. For instance, Cas12k collaborates with Tn7-like transposases for RNA-guided site-specific transposition^18^, whereas Cas12f adopts a unique homodimer architecture crucial for DNA targeting^19^. Conversely, Cas12m diverges from the traditional DNA cleavage paradigm, opting instead for a gene silencing approach to counter MGE threats^20^.

Our investigations into genomic repositories have unearthed a collection of hitherto unidentified Cas12 variants. A notable observation within certain proteins includes regions characterized by low confidence scores in AlphaFold’s structural predictions, which may indicate the presence of Intrinsically Disordered Regions (IDRs)^21–23^. Unlike structured protein domains, IDRs exhibit a high degree of conformational flexibility, allowing them to adopt multiple conformations and interact with a wide range of binding partners. This adaptability enables IDRs to participate in a variety of protein- protein interaction (PPI) networks, mediating complex regulatory mechanisms and signaling pathways within the cell^24,25^.

The large-scale and high-throughput PPIs analyses of the Cas12 variants with potential IDRs uncovered multiple possible interactions, wherein the bacterial thioredoxin TrxA forms a functional heterodimer with a phage-derived entity, referred to here as the Cas12 hacker, enhancing the DNA cleavage capacity and CRISPR immunity of the Cas12 hacker. This mechanism suggests a scenario in which bacteriophages may exploit bacterial components to counteract competing MGEs. This discovery offers important insights into the nuanced coevolutionary ballet between bacterial hosts and their viral aggressors, and proposes novel avenues to refine CRISPR-Cas-based gene editing technologies for enhanced performance and precision.

## Results

### Mining of novel type V CRISPR-Cas12 systems

To systematically explore the diversity within the Cas12 family, we developed a robust and efficient method for identifying novel type V miniature CRISPR-Cas12 systems (Fig. 1a). First, we used CRISPR arrays as reference points to identify adjacent target regions. We then examined open reading frames (ORFs) neighboring the CRISPR arrays to detect potential miniature Cas12 proteins. Following their identification, we classified these proteins, setting aside previously known subtypes to ensure a focus on novel entities. To improve the precision of our classification, we integrated AlphaFold2 prediction for structural analysis of the potential Cas12 proteins (Fig. 1b).

**Fig. 1.**
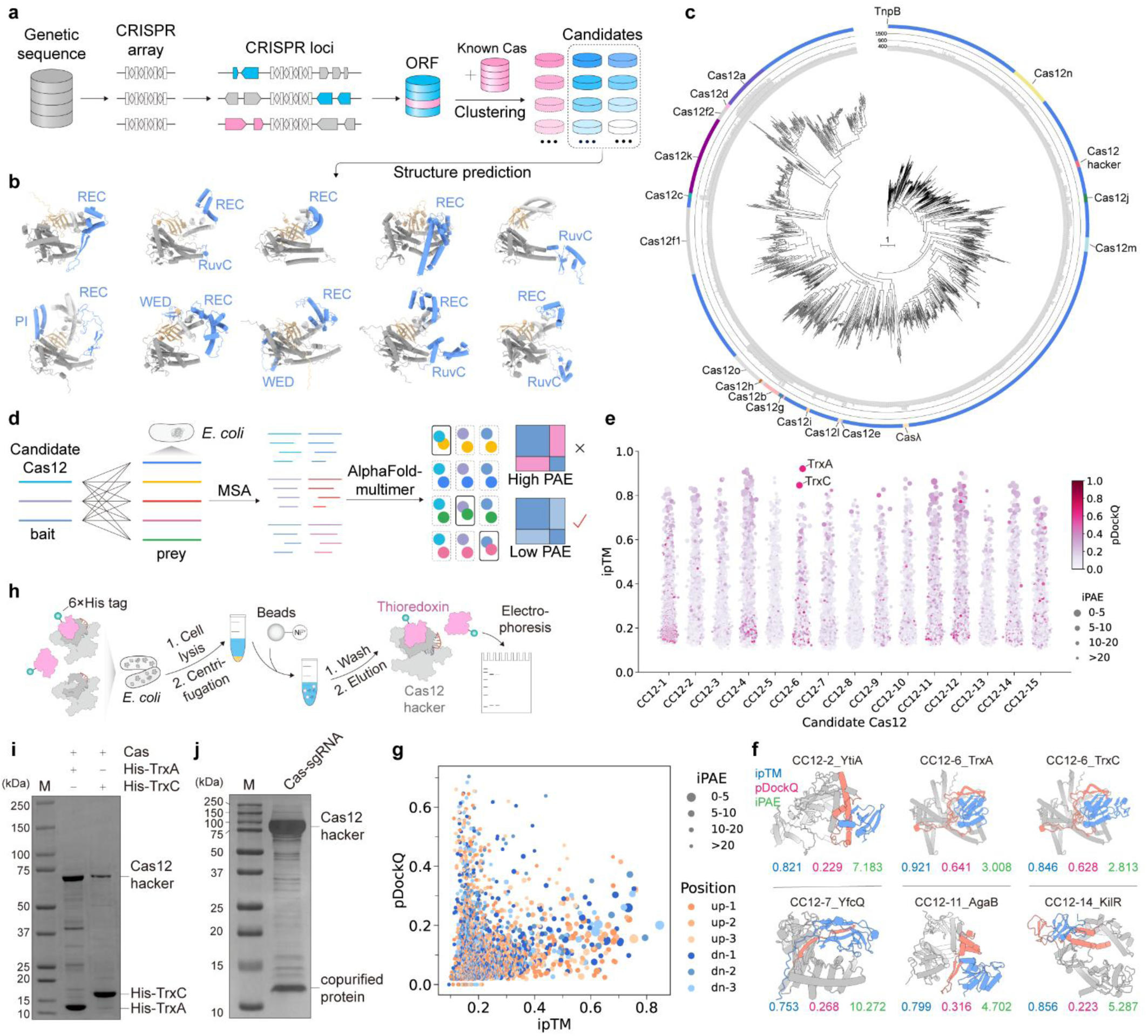
Mining and exploration of type V CRISPR-Cas12 systems. **a**, Overview of the bioinformatics pipeline utilized for mining type V CRISPR-Cas12 systems. **b**, Structural comparison of some selected Cas12 candidates, showcasing the heterogeneity in structural features. Predictive structural models were generated via AlphaFold2, the expanded domains are highlighted in blue. **c**, Phylogenetic analysis of TnpB and Cas12 protein families. Candidate Cas12 (CC12) variants are denoted in blue on the outer ring, while TnpB and characterized Cas12 proteins are color-coded by distinct family groups. Protein lengths are shown in gray bars on the inner ring. **d**, Schematic of AI-driven high-throughput protein-protein interaction (PPI) screening between CC12 proteins and the *Escherichia coli* (*E*. *coli*) proteome. **e**, Bubble plot of the PPI screening results. The predicted complex structures are evaluated using ipTM, pDockQ, and iPAE as scoring metrics. The X-axis represents different CC12 proteins and the Y-axis denotes the ipTM scores. A color gradient ranging from purple to red serves as the legend for the ipTM values, varying dot sizes provides a legend for iPAE values. Notably, the complex structures of CC12-6 (Cas12 hacker) with TrxA and TrxC got the highest scores when assessed using all three metrics simultaneously. **f**, Structural predictions of some high-scoring binary complex by AlphaFold-Multimer. The CC12 proteins, *E*. *coli* proteins, and the potential interaction interfaces of the complexes are represented in gray, blue, and orange, respectively. The values of ipTM, pDockQ, and iPAE are shown in blue, violet, and green respectively. **g**, Bubble plot visualized the PPIs prediction results of CC12 proteins and neighboring open reading frames (ORFs). The scoring metrics for PPIs screening include ipTM, pDockQ, and iPAE. The X-axis, the Y-axis, and varing dot size represent ipTM, pDockQ, and iPAE, respectively. Orange dots represent upstream ORFs, and blue dots represent downstream ORFs. ‘up-1’, ‘up-2’, and ‘up-3’ represent the first, second, and third upstream ORF, respectively; ‘dn-1’, ‘dn-2’, and ‘dn-3’ represent the first, second, and third downstream ORF, respectively. **h**, Schematic of the His tag pull-down assay using untagged Cas12 hacker protein and His-tagged thioredoxin in *E*. *coli*. CC12-6 and thioredoxin are indicated in grey and violet, respectively. **i**, Result of his tag pull-down assay for untagged Cas12 hacker with His-tagged TrxA or TrxC. **j,** SDS-PAGE analysis of sumo_Cas12 hacker-sgRNA RNP purified from *E*. *coli* BL21 (DE3). The molecular weight of co-purified protein is approximately 12 kDa.

This methodological approach enabled us to uncover multiple previously uncharacterized Cas12 families, revealing an unexpected diversity (Fig. 1c). The structural analysis of these Cas12 candidates revealed a common architecture composed of WED, REC, and RuvC domains (Fig. 1b). Despite this overall similarity, we observed notable variations in the size and structure of these domains, especially within the REC domain, suggesting evolutionary adaptations that may confer unique functional traits or roles in specific biological contexts (Fig. 1b).

### Discovery of the host factors associated with Cas12 hacker

Our investigation has revealed that some predicted Cas12 structures may harbor IDRs (Supplementary Fig. 1a), which typically suggest potential interactions with other biomolecules. Given the incomplete nature of many Cas12 sequence contexts within metagenome-assembled genomes (MAGs), establishing their full genomic context proves challenging. A previous study revealed that Cas12k from *Scytonema hofmanni* is known to associate with the *Escherichia coli* (*E*. *coli*) ribosomal protein S15 for target-specific transposition^26^, exemplifying the type of PPIs we aimed to investigate. Inspired by this, we hypothesized that certain proteins capable of interacting with these IDRs may be conserved across various bacterial species. This led us to employ the complete proteome of the classical model organism *E. coli* as potential ‘prey’ for the PPI screening of MAGs-derived Cas12 proteins (Fig. 1d).

We curated fifteen candidate Cas12 (CC12) proteins with potential IDRs as ‘bait’ for PPIs screening and generated a substantial dataset of 65,715 predicted protein complex structures using AlphaFold-Multimer (Fig. 1d-1f, Supplementary Fig. 1b and Supplementary Table 1)^27,28^. To ensure the accuracy of these complex predictions, we applied a comprehensive evaluation framework employing metrics such as interface predicted template modeling (ipTM), pDockQ^29^, and interface predicted aligned error (iPAE), which together offer an in-depth assessment of protein interface characteristics (Fig. 1e, Supplementary Fig. 1b and Supplementary Table 1). We found that these metrics did not strictly correlate for these predicted complex structures, thus making it challenging to accurately assess the quality of the predicted complexes by relying on a single metric. Nevertheless, among these predicted binary PPIs, CC12-6, named as Cas12 hacker, demonstrated consistently high scores in complex with TrxA (UniProt ID: P0AA25) and TrxC (UniProt ID: P0AGG4) proteins across all evaluated criteria (Cas12 hacker and TrxA: ipTM=0.921, pDockQ=0.641, iPAE=3.008; Cas12 hacker and TrxC: ipTM=0.846, pDockQ=0.628, iPAE=2.812), indicating a robust and reliable interaction (Fig. 1e,f, Supplementary Fig. 1b and Supplementary Table 1). Moreover, CC12-2, CC12-7, CC12-11, and CC12-14 were predicated to interact with *E. coli* YtiA, YfcQ, AgaB, and KilR, respectively (Fig. 1f); however, these predictions carry less confidence than the interactions between Cas12 hacker and TrxA or TrxC (Fig. 1f). Given the likelihood that Cas proteins might form functional complexes with neighboring open reading frames (ORFs), we also conducted PPI predictions for candidate Cas proteins along with the upstream and downstream ORFs. However, the results did not reveal particularly reliable protein complex structures (Fig. 1g).

Thereby, we performed further experimental characterizations of the interactions between Cas12 hacker and TrxA/TrxC. Subsequent investigation through bacteria-based pull-down assays, which utilized untagged Cas12 hacker and His-tagged TrxA/TrxC, validated the binding capabilities of Cas12 hacker with both TrxA and TrxC (Fig. 1h,i).

Furthermore, SDS–PAGE analysis of the Cas12 hacker-sgRNA RNP complex purified by His-tagged Cas12 hacker identified a co-purified protein with an approximate mass of ∼12 kDa (TrxA: 11.8 kDa, TrxC: 15.5 kDa) (Fig. 1j). Mass spectrometry (MS) analysis of this RNP complex confirmed *E. coli* TrxA as the predominant candidate for the co-purified protein. (Supplementary Table 2).

### Characterizations of the CRISPR-Cas12 hacker system

Cas12 hacker (Cas12p) is a novel miniature Cas12 family (500-700 amino acids), which appears to be an early evolutionary intermediate from the ancestor TnpB to Cas12 nucleases (Fig. 2a-c). Our investigation uncovered twelve proteins within this family, with eleven exhibiting over 70% sequence identity, indicating a tight genetic conservation (Supplementary Fig. 2a). The scarcity of Cas12 hacker proteins identified may reflect the current limitations in genomic databases. Interestingly, all Cas12 hacker loci were identified within MAGs, and further evaluations using VirSorter 2^30^ and geNomad^31^ pointed towards a phage origin for Cas12 hacker, highlighted by the presence of hallmark viral genes such as phage tail components nearby the CRISPR-Cas12 hacker loci (Supplementary Fig. 2b and Supplementary Table 3). While the bacterial hosts of Cas12 hacker remain to be determined, these findings suggest an interesting phage-associated lineage for this CRISPR- Cas type.

**Fig. 2.**
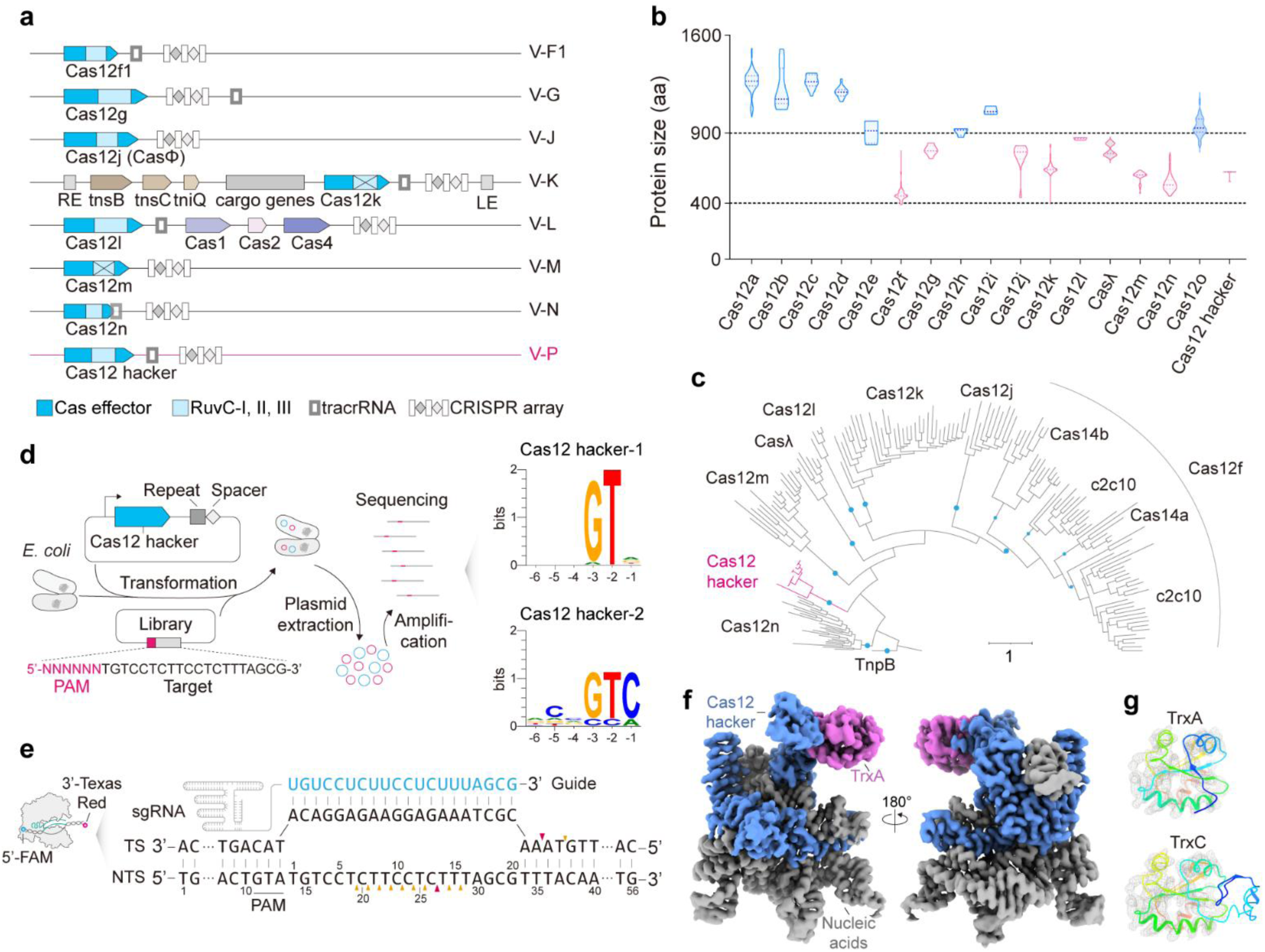
Characterizations of the CRISPR-Cas12 hacker system. **a**, Illustration of representative miniature CRISPR-Cas12 system loci configurations, including V-F1, V-G, V-J, V-K, V-L, V-M, V-N, and V-P. The newly identified CRISPR-Cas12p (Cas12 hacker) system is highlighted in hotpink. **b**, Length distribution of various effector proteins across type V CRISPR-Cas12 systems. The size of Cas12 hacker family effectors ranges from 500 to 700 amino acids. **c**, Phylogenetic tree of the evolutionary relationships between TnpB proteins and representative miniature Cas12 variants, with Cas12 hacker posited as an early evolutionary intermediate. The blue circles represent different Cas12 subtypes. **d**, Schematic of the PAM depletion assay (left) utilized to determine the PAM for the CRISPR-Cas12 hacker systems and webLogo of PAM sequences (right) for Cas12 hacker-1 and Cas12 hacker-2. **e**, Experimental schematic (left) for the cleavage assay using FAM/Texas red-labeled dsDNA substrates with Cas12 hacker and corresponding cleavage patterns (right). Major and minor cleavage products are indicated by red and yellow arrowheads, respectively. **f**, Cryo-EM structural elucidation of the Cas12 hacker-TrxA-sgRNA-target DNA complex, involving 29-nt TS and 11-nt NTS. Key components are colored accordingly: Cas12 hacker protein, nucleic acids, and TrxA are indicated in blue, grey, and violet, respectively. **g**, Structural validation of co-purified protein. TrxA (top) and TrxC (bottom) align well with the globular electron density map. Notably, TrxC encompasses an additional segment absent in TrxA, which lacks corresponding electron density. The structures of TrxA and TrxC were predicted by AlphaFold2 and coloured using rainbow palette, transitioning from the N terminus in blue to the C terminus in red.

We selected Cas12 hacker-1 and Cas12 hacker-2 for initial functional characterization because of their significant disparities in sequence identity and structure (Supplementary Fig. 2a,c). To identify the protospacer adjacent motif (PAM) for Cas12 hacker, we employed a PAM depletion assay that involved transformation of *E. coli* with two plasmids: one encoding the Cas12 hacker-sgRNA complex, the other comprising a library of plasmids with randomized PAM sequences (Fig. 2d). Through next-generation sequencing (NGS), we quantified relative PAM depletions, revealing a favored GTN PAM by both Cas12 hacker nucleases (Fig. 2d and Supplementary Fig. 3a), akin to the preference for ShCas12k^18^. Noteworthy is Cas12 hacker-1’s markedly higher PAM depletion efficiency than Cas12 hacker-2 (Supplementary Fig. 3a), suggesting greater efficacy within *E. coli*. Hence, we centered our subsequent research on Cas12 hacker-1. Consistent with our NGS data, *in vitro* cleavage assays confirmed that Cas12 hacker-1 favored PAM sequences in the order of GTA > GTG > GTT > GTC (Supplementary Fig. 3b). For the sake of brevity, we will refer to ‘Cas12 hacker-1’ as simply ‘Cas12 hacker’ in all future discussions.

To determine the RNA components essential for Cas12 hacker activity, RNA sequencing (RNA-Seq) was conducted, revealing low expression levels of the tracrRNA when heterologously expressed in *E. coli* (Supplementary Fig. 3c). However, a modified RNA co-purified with the Cas12 hacker ribonucleoprotein (RNP) complex confirmed the inclusion of both tracrRNA and crRNA (Supplementary Fig. 3d), suggesting potential RNA degradation within a cellular context.

Investigations into the requisite conditions for effective Cas12 hacker-mediated DNA cleavage identified the critical roles of divalent ions and NaCl in catalysis (Supplementary Fig. 3e). Cas12 hacker exhibited superior activity across a temperature range of 30°C to 40°C and was active in the presence of magnesium or manganese ion (Supplementary Fig. 3e), showcasing a flexibility uncommon in many Cas proteins traditionally reliant on magnesium. In-depth analysis of Cas12 hacker’s cleavage patterns employed urea-PAGE-assisted cleavage assays, revealing its capability to generate primary cuts on both target strand (TS) and non-target strand (NTS), leading to sticky ends distal to the PAM site (Fig. 2e and Supplementary Fig. 3f,g). Progressive degradation of the NTS was observed (Fig. 2e and Supplementary Fig. 3g), indicative of trans-cleavage activity.

### TrxA directly binds to Cas12 hacker

To substantiate the association between Cas12 hacker and *E. coli*-derived TrxA/TrxC, as well as to elucidate the mechanisms underlying Cas12 hacker-mediated DNA cleavage, we co-expressed Cas12 hacker with the corresponding 246-nt sgRNA, composed of a 185-nt tracrRNA, a 4-nt GAAA linker, a 37-nt repeat, and a 20-nt guide in *E*. *coli*. Subsequent purification yielded the Cas12 hacker-sgRNA RNP complex, which was then combined with a 29-nt TS DNA and an 11-nt NTS DNA, comprising a GTA PAM, to form the Cas12 hacker-sgRNA-dsDNA complex (Supplementary Fig. 4a,b). Cryo-electron microscopy (Cryo-EM) analysis facilitated a three-dimensional reconstruction of this complex at an overall resolution of 2.7 Å (Fig. 2f, Supplementary Fig. 4c-h and Supplementary Table 4).

Structural analysis revealed a notable discrepancy: an IDR within the REC domain, as predicted by AlphaFold2, was found to be misaligned with an adjacent globular density (Fig. 2f and Supplementary Fig. 5a). This density, however, exhibited a strong correlation with TrxA (Fig. 2g). Despite the C-terminal domain of TrxC closely resembling the entirety of the TrxA protein, the presence of an additional N-terminal domain in TrxC, which is absent in the cryo-EM map, renders it a less likely match (Fig. 2g). TrxA, with its pronounced presence in the MS results and congruent structure to the cryo- EM density, was identified as the primary host-derived protein integral to the complex (Fig. 2g and Supplementary Table 2).

Thioredoxins (Trx) are ubiquitous proteins involved in various thiol-disulfide redox reactions, sharing a conserved thioredoxin fold and a WCGPC catalytic motif^32^. Moreover, it has been found to interact with a diverse range of protein partners by forming functional complexes crucial for different biological processes. For instance, T7 phage-derived DNA polymerase interacts with *E. coli* TrxA to increase the processivity of nucleotide polymerization^33,34^; *E. coli* 3’- Phosphoadenosine-5’-phosphosulfate (PAPS) reductase^35^, human TXNIP^36^, and NLRP1^37^ also bind to thioredoxin for biological regulation (Supplementary Fig. 5b-e). Given Cas12 hacker’s origin from metagenomic assemblies, definitively pinpointing the original TrxA variant naturally associated with Cas12 hacker in its native host remains challenging. Nevertheless, our findings securely establish that *E. coli* TrxA can robustly associate with Cas12 hacker.

### Structure of the Cas12 hacker-TrxA-sgRNA-target DNA complex

The resolved complex structure is composed of a Cas12 hacker-TrxA protein heterodimer and a sgRNA-DNA heteroduplex (Fig. 3a). In the structure, Cas12 hacker is mainly divided into two lobes: the REC lobe and the nuclease (NUC) lobe (Fig. 3b). The REC lobe, located at the N-terminus, consists of the WED, REC, thioredoxin binding (TB), and PAM interaction (PI) domains (Fig. 3b). The WED domain is composed of a seven-stranded β-barrel (Fig. 3a and Supplementary Fig. 6a). The REC domain, comprising six α helices, is inserted between the β1 and β2 strands of the WED domain (Fig. 3a and Supplementary Fig. 6a). The TB domain, defined by two pairs of parallel β-strands, is positioned inside of the REC domain and could be sequentially divided into two distinctive subregions (Fig. 3a,b and Supplementary Fig. 6a). The N-terminal subregion, termed TB.a, is enriched in hydrophobic amino acids, while the C-terminal counterpart, TB.b, abounds with basic amino acids (Fig. 3b and Supplementary Fig. 6b). The PI domain, flanked by the WED and RuvC domains, consists of two parallel helices and is similar to that found in Cas12k (Fig. 3a and Supplementary Fig. 6a)^26,38^. The NUC lobe, located at the C-terminus, consists of the RuvC and zinc finger (ZF) domains (Fig. 3b). The RuvC domain exhibits an RNase H fold, with residues D378, E478, and D582 constituting a catalytic center (Fig. 3a and Supplementary Fig. 6a,c). The ZF domain is inserted between the β5 strand and the α5 helix (Fig. 3a and Supplementary Fig. 6a). The C-terminal domain (CTD), likely to be an evolutionary legacy of TnpB proteins, was disordered in the structure and exhibited little sequence similarity to other Cas12 proteins (Fig. 3a,b). Overall, the structural characteristics of Cas12 hacker proteins display notable similarities with TnpB and Cas12k, particularly within the WED and RuvC domains. However, a distinctive difference is observed in the REC domains of Cas12 hacker and Cas12k, which are substantially larger than the more compact REC domain seen in TnpB (Supplementary Fig. 6a)^39^. This size variation in the REC domains underscores their enhanced capacity for interactions with other partner proteins. Moreover, structural comparisons of TnpB and Cas12 nucleases highlight significant diversification, which makes Cas12 proteins with remarkable plasticity and expansiveness in function, although they all share similar domain configurations to TnpB (Supplementary Figs. 7 and 8)^26,39–53^.

**Fig. 3.**
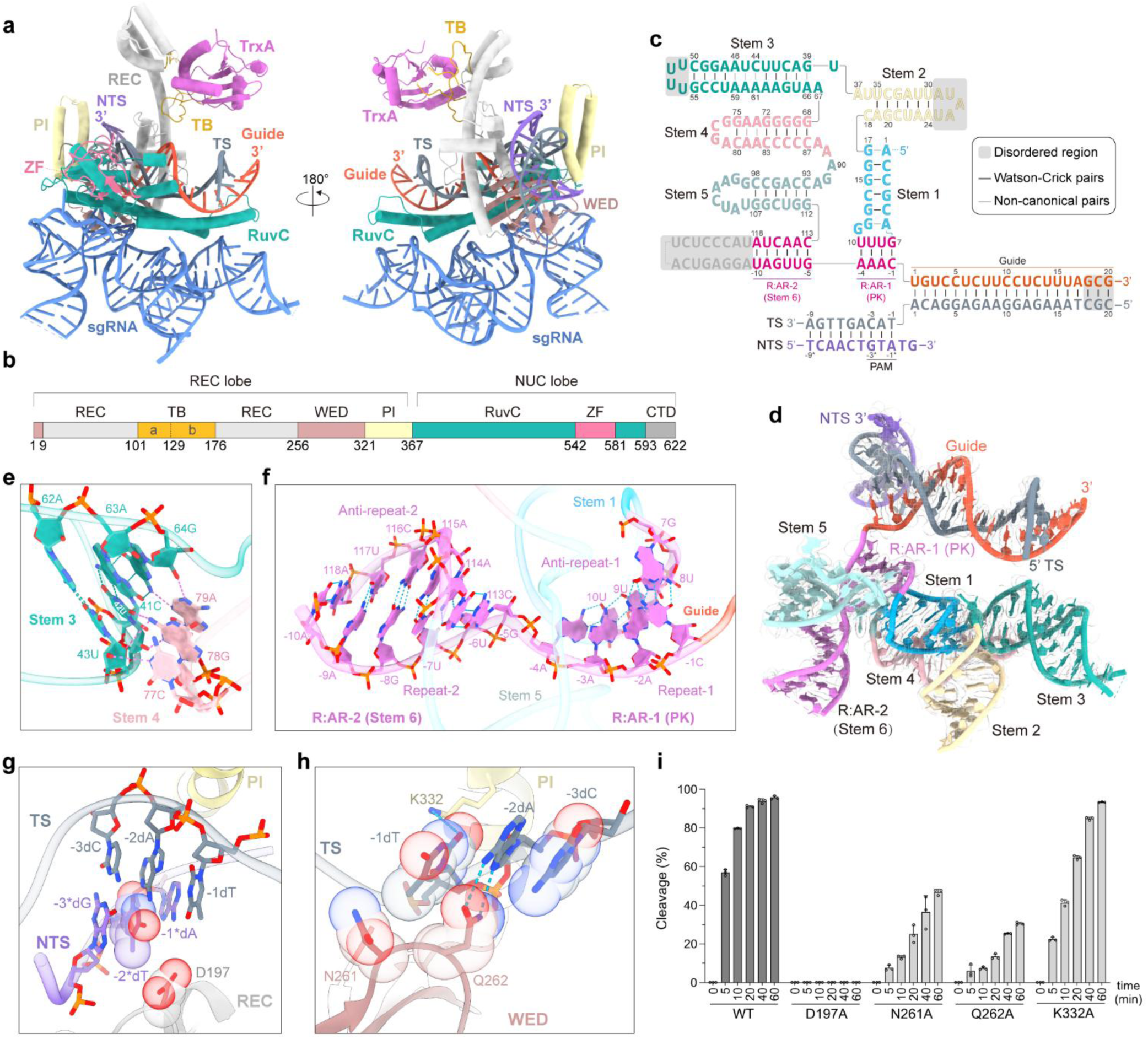
Cryo-EM structure of the Cas12 hacker-TrxA-sgRNA-target DNA complex. **a**, The overall atomic model of the Cas12 hacker-TrxA-sgRNA-target DNA (29-nt TS and 11-nt NTS) complex. **b**, Domain organization of Cas12 hacker. **c**, Schematic of sgRNA and target DNA. **d**, Atomic model of sgRNA and target DNA. **e**, Close-up view of the stem3-stem4 interaction. Hydrogen bonds are indicated by dashed lines. **f**, Close-up view of the crRNA repeat:tracrRNA anti-repeat (R:AR) duplex, highlighting hydrogen bonds with dashed lines. **g**,**h**, Zoom-in views illustrating the base-specific interactions of Cas12 hacker with the NTS (**g**) and TS (**h**) within the PAM region of the DNA duplex. Hydrogen bonds are shown with dashed lines, while van der Waals interactions are represented by space-filling models. **i,** Comparison of *in vitro* cleavage activities between the wild-type (WT) Cas12 hacker and mutants deficient in PAM recognition (D197A, N261A, Q262A, and K332A). Data are represented as mean ± SD (n = 3).

The sgRNA of Cas12 hacker consists of a 118-nt tracrRNA and a 27-nt crRNA, which includes a 10-nt repeat and a 17-nt guide sequence (Fig. 3c). This complex structure encompasses six stems and a pseudoknot (PK) (Fig. 3c,d). The stem 4 interacts with the stem 3 through nucleobases of 77C, 78G, and 79A, which engage in hydrogen-bonding interactions with 43U, 42U, and 64G (Fig. 3e). The PK arises from base pairing between crRNA -1C to -4A and tracrRNA 7G to 10U, and is referred to as repeat-anti-repeat duplex 1 (R:AR-1) (Fig. 3c,f). Consistent across the CRISPR-Cas12 hacker family, multiple sequence alignments highlight a conserved AAAC: UUUG base pairing motif for PK formation (Supplementary Fig. 9a). Notably, despite diversity in sequence and structure, tracrRNAs of Cas12 enzymes characteristically engage in PK formation with the 3’ ends of the repeat sequences (Supplementary Fig. 9b)^26,39,41–44,48^, underscoring the PK’s critical role in enzyme functionality. Further examination unveiled that the RNA architecture of Cas12 hacker demonstrates both similarities and divergences when compared to those of TnpB and Cas12k (Supplementary Fig. 9c)^26,39^. Specially, stems 1∼4 and 6 of Cas12 hacker’s sgRNA exhibit structural consonance with the stems of TnpB’s ωRNA (Supplementary Fig. 9c)^39^. In contrast, the sgRNA of Cas12k presents a more complex structure, bridging Cas12k to Tn7-like transposases (Supplementary Fig. 9c,d)^26^. In the interactions between Cas12 hacker and sgRNA, the stems 3 and 5 of sgRNA, as well as its R:AR-1 region, engage with the WED and RuvC domains through hydrogen bonds and electrostatic forces (Supplementary Figs. 10 and 11a-d). The sgRNA-target DNA heteroduplex is accommodated by the positively charged channel formed by the REC, WED and RuvC domains (Supplementary Figs. 10 and 11a,e,f).

Moreover, our exploration extended to the PAM recognition mechanism, revealing a collaborative role of the WED, REC, and PI domains in securely engaging the GTA PAM-containing DNA duplex (Fig. 3g,h and Supplementary Fig. 11g). The interactions involve the -2*dT nucleobase on the NTS, which connects with D197 within the REC domain through van der Waals forces (Fig. 3g). Correspondingly, on the TS, the -1dT nucleobase establishes a hydrogen bond with K332 in the PI domain, further engaging N261 and Q262 in the WED domain via van der Waals interactions (Fig. 3h). In addition, the -2dA nucleobase forms a hydrogen bond, while the -3dC nucleobase interacts with Q262’s side chain through van der Waals forces (Fig. 3h). Our mutational analysis highlights the important nature of these interactions. Specifically, introducing the D197A mutation completely abolishes the cleavage ability of Cas12 hacker (Fig. 3i). The N261A and Q262A mutations significantly impair its activity, and the K332A mutation results in a slight decrease in its enzymatic performance (Fig. 3i). Despite significant sequence divergence, structural analyses highlight a shared PI domain layout between Cas12 hacker and Cas12k, albeit with a notable 180-degree rotational disparity in Cas12k’s PI domain relative to Cas12 hacker’s (Supplementary Fig. 12a-d). This rotation alters the helical contacts with the PAM-containing DNA, yet the vicinity near the PAM site in each PI domain maintains a composition rich in basic residues (Supplementary Fig. 12e-g).

### Molecular interactions between TrxA and Cas12 hacker

To gain insight into the molecular mechanisms of how TrxA associates with Cas12 hacker, we analyzed the detailed interactions between TrxA and Cas12 hacker. Our analysis revealed that the interaction is chiefly governed by hydrophobic forces (Fig. 4a-c and Supplementary Fig. 13a). The redox center of TrxA, key to its function and known to facilitate interactions with various proteins^35–37^, is surrounded by a cluster of hydrophobic residues (W32, P35, M38, I61, and I76) (Fig. 4b,c and Supplementary Fig. 13a). This redox center forms a stable disulfide bond between C33 and C36 (Fig. 4b and Supplementary Fig. 13a,b).

**Fig. 4.**
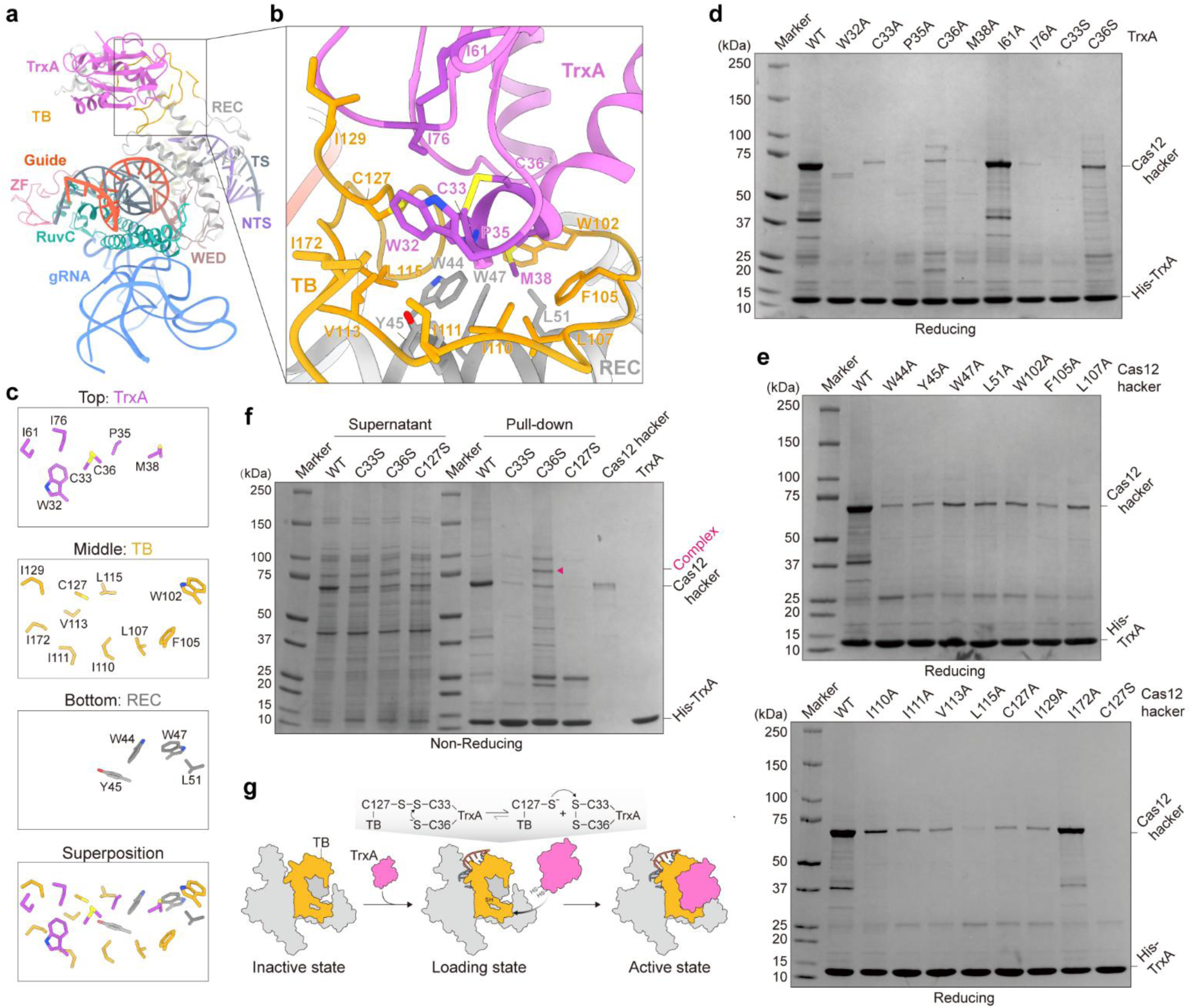
Molecular insights into TrxA association with Cas12 hacker. **a**, The overall atomic model of the Cas12 hacker-TrxA-sgRNA-target DNA (29-nt TS and 11-nt NTS) complex. **b**, Close-up view of the detailed interactions between the TB domain and TrxA. TrxA forms a C33-C36 intramolecular disulfide bond. **c**, Depiction of key residues within the three-tiered hydrophobic interface. **d**, SDS-PAGE results from pull-down assays comparing the interaction between wild-type (WT) Cas12 hacker and wild-type or mutant His-TrxA, visualized with CBB staining. **e**, SDS-PAGE analysis of pull-down assays investigating the binding between wild-type or mutant Cas12 hacker proteins and wild-type His-TrxA, visualized with CBB staining. **f**, Analysis of intermolecular disulfide bond formation in wild-type Cas12 hacker and wild-type or mutant His-TrxA, as well as Cas12 hacker mutant (C127S) with wild-type His-TrxA using non-reducing SDS-PAGE, visualized with CBB staining. **g**, A proposed model illustrating the assembly process between Cas12 hacker and TrxA protein, emphasizing the transformation of intermolecular to intramolecular disulfide bonds.

The corresponding hydrophobic interface on Cas12 hacker comprises residues W102, F105, L107, I110, I111, V113, L115, C127, I129, and I172 within the TB domain (Fig. 4b,c), along with W44, Y45, W47, and L51 from the REC domain (Fig. 4b,c). Together, they create a hydrophobic pocket that snugly accommodates TrxA’s redox center (Fig. 4b,c and Supplementary Fig. 13a). This interaction interface is remarkably ordered, with the TrxA redox center at the top, the TB domain of Cas12 hacker forming the middle layer, and the Cas12 hacker REC domain constituting the bottom layer, exemplifying a tri-layered hydrophobic docking structure (Fig. 4c and Supplementary Fig. 13a).

To validate the roles of those residues in Cas12 hacker-TrxA association, we conducted pull-down and bacterial genome interference assays using the Cas12 hacker-TrxA interface mutants. Disruption of the binding site through alanine substitution of TrxA W32, P35, M38, and I76, the side chain of which was inserted into the Cas12 hacker hydrophobic pocket, nearly abolished the binding and genome interference activity of Cas12 hacker (Fig. 4b,d and Supplementary Fig. 13a,c,d). Subsequently, we assessed the impact of residues within the hydrophobic pocket of Cas12 hacker. Consistent with the structural analysis, mutations in the REC domain (W44A, Y45A, W47A, and L51A) and the TB.a domain (W102A, F105A, L107A, I110A, I111A, V113A, L115A, C127A, and I129A) drastically hindered TrxA binding and substantially compromised the genome interference activity of Cas12 hacker (Fig. 4b,c,e and Supplementary Fig. 13e,f). These results demonstrated that the hydrophobic pocket formed by TB and REC domains is vital for TrxA association.

The close proximity of TrxA C33/C36 and Cas12 hacker C127 emphasizes the likelihood that redox dynamics play a crucial role in their interaction. Cryo-EM studies revealed a noticeable density between C33 and C36 of TrxA, indicating a stable formation of the C33-C36 disulfide bond (Supplementary Fig. 13b). Subsequent experiments utilizing non-reducing SDS-PAGE showed that the wild-type Cas12 hacker could form an intermolecular disulfide bond with the TrxA (C36S) mutant, a phenomenon not observed with either the wild-type or the TrxA (C33S) mutant (Fig. 4f). In contrast, the Cas12 hacker (C127S) mutant was unable to form an intermolecular disulfide bond with the wild-type TrxA (Fig. 4f), underscoring the specificity of this cysteine-dependent interaction.

Furthermore, the intermolecular disulfide bond between the wild-type Cas12 hacker and the TrxA (C36S) mutant was significantly diminished when subjected to reducing SDS-PAGE (Supplementary Fig. 13g). Additionally, mutating either TrxA C33, C36, or Cas12 hacker C127 to alanine or serine differentially reduced the binding between Cas12 hacker and TrxA (Fig. 4d,e and Supplementary Fig. 13c,e). Notably, mutations at TrxA C33 and Cas12 hacker C127, but not at TrxA C36, significantly impaired the interference activity (Supplementary Fig. 13d,f), suggesting that altering TrxA C36 might facilitate the formation of the intermolecular disulfide bond, thereby preserving Cas12 hacker functionality.

Based on these observations, we hypothesize that the interaction mechanism includes a temporary intermolecular disulfide bond between Cas12 hacker C127 and TrxA C33. This bond is then reduced by the nearby C36, leading to a stable C33-C36 disulfide bond within TrxA (Fig. 4g). This redox-driven mechanism is crucial for the precise and efficient binding needed for the augmented nuclease activity of Cas12 hacker. A similar interaction involving human NLRP1 and thioredoxin demonstrates the relevance of this mechanism (Supplementary Fig. 5e)^37^.

### TrxA binding activates Cas12 hacker’s DNA cleavage capacity

Given the robust binding between Cas12 hacker and TrxA, we delved into TrxA’s influence on Cas12 hacker-mediated DNA cleavage through comprehensive bacteria-based genome interference and *in vitro* DNA cleavage assays. Genome targeting by Cas12 hacker significantly hindered the growth of the wild-type strain (Fig. 5a,b). Intriguingly, the disruption of the *trxA* gene markedly diminished Cas12 hacker’s interference efficacy, which was rescuable upon the reintroduction of the wild-type *trxA* gene (Fig. 5c). Additionally, purifying Cas12 hacker RNP from the *E. coli trxA*-knockout strain proved challenging, resulting in substantially reduced DNA cleavage activity compared to the wild-type strain (Fig. 5d). Supplementing TrxA into the assay system led to a partial recovery of Cas12 hacker cleavage activity, yet optimal activity eluded attainment even with an excess of TrxA (Supplementary Fig. 14a). Furthermore, both bacteria-based genome interference and *in vitro* DNA cleavage assays underscored the minimal effect of another thioredoxin, TrxC, on Cas12 hacker’s function, corroborating the specificity of TrxA as Cas12 hacker’s primary interaction partner (Fig. 5b,d).

**Fig. 5.**
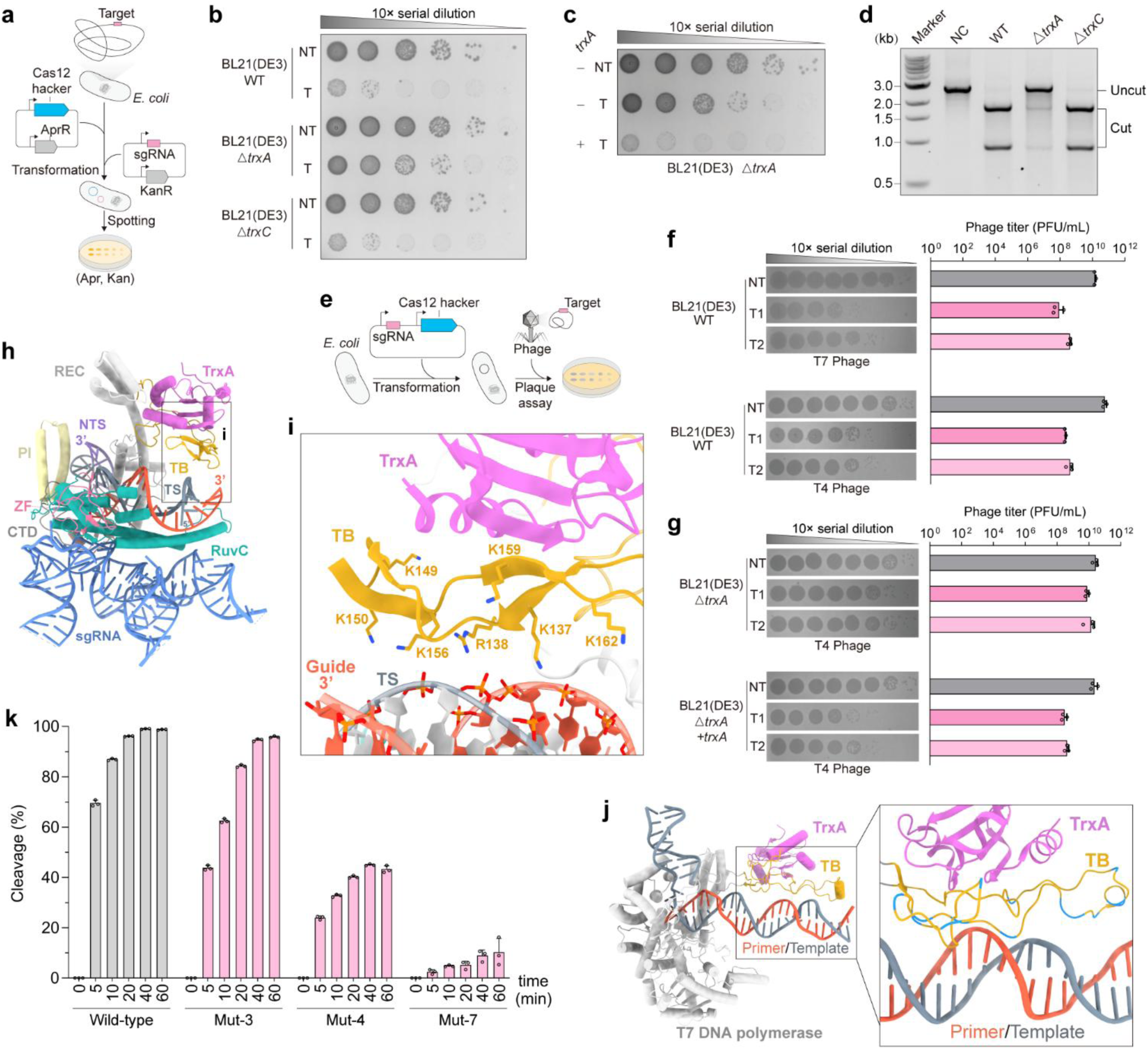
TrxA is essential for Cas12 hacker’s DNA cleavage activity. **a**, Schematic of *E*. *coli* genome interference assay using CRISPR-Cas12 hacker. **b**, Spot dilution assays evaluating CRISPR-Cas12 hacker functionality in *E. coli*, comparing wild-type (WT), Δ*trxA*, and Δ*trxC* strains with a non-targeting (NT) or targeting (T) sgRNA. **c**, Spot dilution assays assessing CRISPR-Cas12 hacker activity in Δ*trxA E. coli* with (+) or without (–) *trxA* complementation. **d**, *In vitro* DNA cleavage assays with Cas12 hacker proteins purified from wild-type, Δ*trxA*, or Δ*trxC E. coli* strains. NC indicates negative control. **e**, Schematic for phage plaque assay using CRISPR-Cas12 hacker. **f**, Phage plaque assays of T4 and T7 phages on *E. coli* BL21 (DE3) harboring CRISPR-Cas12 hacker with non-targeting (NT) or targeting (T) sgRNA. Results for plaque-forming units (PFUs) per milliliter are presented as mean ± SD (n = 3). **g**, T4 phage plaque assays on Δ*trxA E. coli* BL21 (DE3) with (+*trxA*) or without *trxA* complementation. PFU/mL data are displayed as mean ± SD (n = 3). **h**, Overall structure of the Cas12 hacker-TrxA-sgRNA-target DNA (33-bp dsDNA) complex. **i,** Close-up view of the interaction between the Cas12 hacker TB domain and the DNA-RNA duplex. **j.** Structure of T7 DNA polymerase in complex with primer-template DNA duplex. Lysine and arginine residues are indicated in blue. **k**, Comparative *in vitro* cleavage assays for wild-type and mutant Cas12 hacker proteins. Mut-3 represents a triple mutant K149A, K150A, and K156A, while Mut-4 is a quadruple mutant K137A, R138A, K159A, and K162A, and Mut-7 is a septuple mutant combining mutations from Mut-3 and Mut-4. Data presented as mean ± SD (n = 3).

Expanding upon TrxA’s ubiquity and evolutionary conservation, we explored its function with Cas12 hacker in other bacterial species, specifically *Klebsiella pneumoniae* 1.6366 and *Acinetobacter baumannii* 17978. Genome interference assays confirmed that intrinsic TrxA within these strains is similarly essential for activating Cas12 hacker (Supplementary Fig. 14b,c), hinting at a widespread applicability between Cas12 hacker and TrxA across diverse bacterial hosts. It is worth noted that deletion of *trxA* markedly hindered *A. baumannii* growth (Supplementary Fig. 14c).

We further assessed the biological implications of Cas12 hacker-TrxA association through phage plaque assays. In wild-type *E. coli*, the CRISPR-Cas12 hacker system conferred protection against both T4 and T7 phages by targeting their genomes, significantly reducing phage plaque formation (Fig. 5e,f). However, in *trxA*-knockout strains, T4 phage infections increase notably (Fig. 5g), while T7 phage infections hindered (Supplementary Fig. 14d), likely owing to TrxA’s indispensable role in T7 DNA polymerase function, fundamental for T7 phage DNA replication^54^. Reintroducing *trxA* into the knockout strain restored the CRISPR-Cas12 hacker system’s protective capabilities against T4 phage challenges (Fig. 5g). These findings spotlight Cas12 hacker’s synergy with host TrxA protein as a multifaceted approach to combat foreign DNA threats, showcasing the intricate defense strategies unleashed by the CRISPR-Cas12 hacker system in bacterial immunity.

### Molecular mechanism of Cas12 hacker activation by TrxA

In the initial cryo-EM map, an intact density for the TB.b loop was conspicuously absent (Supplementary Fig. 5a), likely due to its flexibility, which presented a challenge for elucidating the mechanistic basis of TrxA’s associative activation of Cas12 hacker. To better decode the molecular mechanisms, we conducted refined cryo-EM studies with an extended 33 bp double-stranded DNA (dsDNA), hypothesizing that the longer NTS might stabilize the TB.b loop. Advanced 3D classification techniques subsequently unveiled a novel conformation with the TB.b’s flexible loop proximal to the PAM- distal terminus of the sgRNA-target DNA heteroduplex, leading us to determine a structure at a resolution of 3.2 Å (Fig. 5h, Supplementary Figs. 15a-g and 16a,b and Supplementary Table 4).

An analysis of the TB.b loop revealed an abundance of positive surface charges that engaged favorably with the negatively charged sgRNA-DNA duplex backbone (Fig. 5h,i and Supplementary Fig. 16c,d), mirroring the DNA-binding mechanism of T7 DNA polymerase’s TB domain,^33,55^ which employs basic amino acids to enhance nucleic acid affinity (Fig. 5j and Supplementary Fig. 16e). Significantly, an analysis of the interface between TrxA and the TB flexible loop in both Cas12 hacker and T7 DNA polymerase uncovers the presence of hydrophobic amino acids, implicating non-polar interactions as the primary facilitators for the movement of the loop (Supplementary Fig. 16c-f). Furthermore, the anchoring of TrxA likely serves to restrict the TB flexible loop’s range of movement, conferring spatial precision and precluding non-specific motion (Fig. 5h,i,j and Supplementary Fig. 16b,c,e).

To further probe the role of these charged residues in TB.b domain, we performed alanine substitutions at lysine and arginine sites (K137, R138, K149, K150, K156, K159, and K162) adjacently positioned to the RNA-DNA duplex (Fig. 5i and Supplementary Fig. 16c,d). *In vitro* DNA cleavage assays revealed that individual residue mutations had negligible effects on Cas12 hacker’s nuclease activity (Supplementary Fig. 16g). Nonetheless, we observed that concurrent mutations at either end of the loop, namely K149A, K150A, and K156A, led to a modest decline in activity, whereas alterations situated centrally, including K137A, R138A, K156A, and K159A, elicited a significant reduction in cleavage efficiency (Fig. 5k). When all seven basic residues were mutated simultaneously, a near-complete cessation of DNA cleavage activity was noted (Fig. 5k), highlighting the indispensable contribution of these charged residues in Cas12 hacker functionality. These observations underscore the pivotal role of TrxA in fostering interactions between the dynamic TB.b loop and nucleic acids, thereby potentiating the DNA cleavage capacity of Cas12 hacker.

## Discussion

In this study, through metagenomic exploration and high-throughput PPIs analyses, we have discovered a phage-encoded, miniature Cas12 hacker nuclease, which intriguingly partners with the bacterial host’s thioredoxin protein, TrxA. This partnership not only proves critical for the activation and genome interference efficiency of the Cas12 hacker nuclease but also uncovers a previously unexplored aspect of CRISPR immunity. Unlike the conventional focus on CRISPR inhibition mechanisms, our findings illustrate how host factors can be leveraged to augment CRISPR activity (Fig. 6a). Such synergy exemplifies an evolutionary novelty where phages co-opt host components to bolster their CRISPR arsenal against competitive MGEs.

**Fig. 6.**
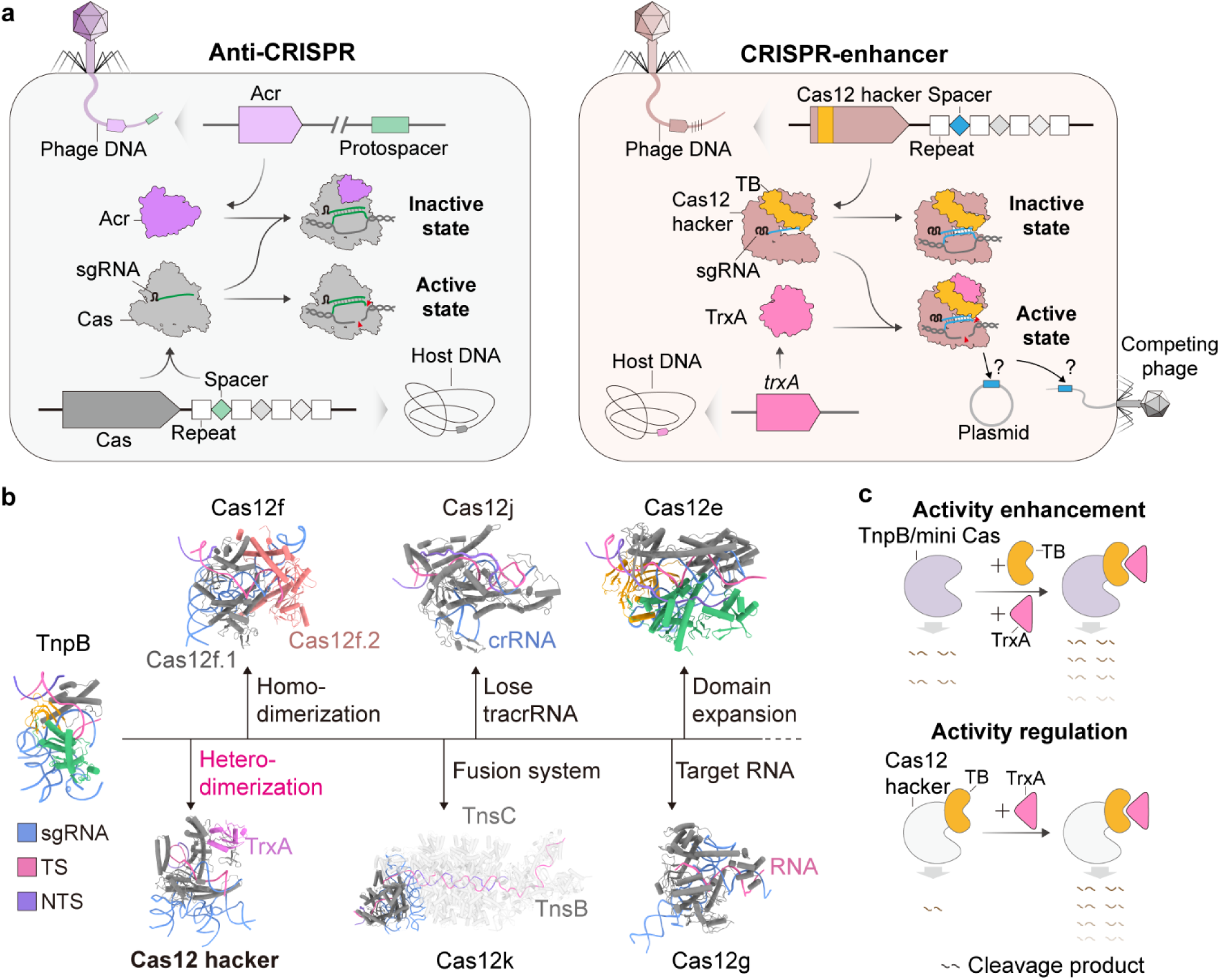
Schematic representation for CRISPR evolutionary mechanisms and pathways. **a**, Conceptual illustration presenting the mechanistic model of Anti-CRISPR (Acr) and CRISPR-enhancer systems. **b**, Evolutionary diagram tracing the structural and functional progression of CRISPR-Cas12 systems, from early archetypal architectures to advanced derivatives. The following representative systems are included: TnpB (PDB ID: 8H1J), Cas12f (PDB ID: 8J12), Cas12j (PDB ID: 7LYT), Cas12e (PDB ID: 7WAY), Cas12g (PDB ID: 6XMG), Cas12k (PDB ID: 8EA4), and Cas12 hacker. **c,** Potential application prospects for TrxA as a CRISPR-enhancer.

Through integrative approaches, including cryo-EM, biochemical, bacterial genome interference, and phage plague assays, we established TrxA as a pivotal cofactor that activates Cas12 hacker’s nuclease activity by forming a distinctive heterodimer structure. This mode of activation starkly contrasts with the diversification strategies observed in the wider Cas12 family, which vary from protein domain expansion (large Cas12 nucleases) and homodimerization (Cas12f) to protein polymerization associated with a transposition system (Cas12k)^17^, thus emphasizing the myriad evolutionary paths Cas nucleases have traversed (Fig. 6b).

A detailed structural examination of the Cas12 hacker-TrxA interplay highlights the essential role of the TB domain in facilitating TrxA recruitment. Mutagenesis studies have elucidated that the TB.a and REC domains form a hydrophobic pocket to effectively capture TrxA, analogous to the interaction between NLRP1 and thioredoxin^37^. Furthermore, TB.b- nucleic acids interaction revealed a complex mechanism paralleled by the T7 phage-derived DNA polymerase^54,55^, underscoring the broad strategy of harnessing host factors to enhance the enzymatic efficiency of phage-encoded proteins. Miniature Cas12 nucleases, along with their ancestral TnpB nucleases, have emerged as promising genome editing tools due to their compact size and potent genome interference capabilities, ideal for delivery with size-constrained vectors^56–58^. However, the inherent genome editing activity of these naturally derived nucleases in human cells is minimal, posing a significant barrier to their application. Through targeted protein engineering and sgRNA optimization, their genome editing efficiency can be substantially improved^59,60^. Building on these strategies, our discovery that host factor TrxA can act as a CRISPR enhancer presents an innovative method to amplify the activity of miniature Cas nucleases without notably increasing their molecular size by engineering the TB domain into these mini nucleases (Fig. 6c).

Additionally, because the Cas12 hacker nuclease shows negligible activity without TrxA, this mechanism can be potentially utilized for safer editing by modulating the activity of the Cas12 hacker through TrxA regulation (Fig. 6c).

Collectively, our exploration into the CRISPR hacker system enhances our comprehension of CRISPR immunity, bringing to light the dynamic and cooperative potential within the host-phage evolutionary contest. Such insights not only elucidate the adaptive strategies employed by both bacteria and phages but also pave the way for the development of sophisticated CRISPR-Cas tools, offering enhanced activity and control for a wide array of applications.

## Method Details

### Computational pipeline for analysis of candidate CRISPR-Cas12 systems

For the mining of potential candidate Cas12 proteins, a genomic database was constructed by combining publicly available data from NCBI and JGI. CRISPR arrays were identified using MinCED^61^. Protein-coding genes within 20 kb windows around the CRISPR arrays were predicted using Prodigal^62^. Known Cas12 sequences were collected from previous studies^15,16,63^ and aligned using MAFFT v7^64^. The alignment was converted into a Hidden Markov Model (HMM) profile. The profile was used to search the protein database by HMMER^65^. Candidate Cas12 proteins ranging from 400 to 900 amino acids were selected. These candidates and known Cas12 proteins were combined and clustered using MMSeqs2^66^. The clusters corresponding to known Cas12 families were excluded. Finally, the structures of candidate Cas12 proteins were predicted using AlphaFold2^67,68^.

For PPI screening, fifteen candidate Cas12 proteins containing potential IDRs were curated as bait proteins, and 4,381 proteins from *E*. *coli* were used as potential prey proteins. Bait-prey protein interactions were predicted by AlphaFold- Multimer^27,69^. pDockQ scores were calculated by FoldDock^29^.

For the phylogenetic analysis presented in Fig. 1c, candidate Cas12 proteins were clustered by MMSeqs2, the representative candidates, TnpB, and known Cas12 proteins were aligned by MUSCLE v5^70^ (Super5 algorithm). the phylogenetic tree was constructed by FastTree^71^ (LG+Gamma20). For the phylogenetic analysis presented in Fig. 2c, the sequences of TnpB, Cas12f, Cas12j, Cas12k, Cas12l, Cas12m, Cas12n, Casλ and Cas12 hacker were combined and aligned by MUSCLE v5^70^. The phylogenetic tree was inferred by IQ-TREE v2^72^ using automatic model selection and 1,000 bootstraps. The trees were visualized by iTOL^73^.

### Plasmid construction

The gene locus encoding the CRISPR-Cas12 hacker system was synthesized by Sangon Biotech (Shanghai, China). The *trxA* gene and *trxC* gene were amplified from genomic DNA of *E*. *coli* BL21(DE3). For the PAM depletion assay, the CRISPR-Cas12 hacker locus with the *rpsL* promoter (P*rpsL*) was inserted into a p15A-based vector by Gibson assembly. A plasmid DNA library containing 6-bp randomized PAMs at the 5’ end of the target site was constructed as the procedure previously described^74^. For the bacterial genome interference assay in *E. coli* and *K*. *pneumoniae*, a Cas12 hacker expression cassette was cloned into a p15A-based vector, while a sgRNA_v2-expressing cassette with *Bsa*I sites for target insertion was cloned into a pUC19-based vector via Gibson assembly. The *trxA*-harboring plasmid was constructed by inserting *trxA* coding gene, along with its native promoter into the sgRNA_v2-harboring plasmid. For the genome interference assay in *A*. *baumannii* strains, the Cas12 hacker-harboring plasmid was constructed by replacing the Cas9 gene in the pCasAb plasmid with the Cas12 hacker gene and removing the RecAB recombination system. The sgRNA- harboring plasmid was constructed by replacing the sgRNA-expressing cassette in the pSGAb plasmid with a sgRNA_v2- expressing cassette of Cas12 hacker. For deletion of *trxA* or *trxC* in *E*. *coli* and *K*. *pneumoniae* strains, the pCasKP-pSGKP genome editing system was employed^75^. Specially, the pSGKP-based plasmid was constructed by inserting *trxA*-targeting or *trxC*-targeting spacer via Golden Gate assembly and corresponding repair template via Gibson assembly. For deletion of *trxA* in *A*. *baumannii*, the pCasAb-pSGAb genome editing system was employed^76^. Specially, the pSGAb-based plasmid was constructed by inserting *trxA*-targeting spacer via Golden Gate assembly and corresponding repair template via Gibson assembly. For the phage plaque assay, a P*rpsL*-driven Cas12 hacker nuclease and a J23119-driven sgRNA_v2 were cloned into a p15A-based vector via Gibson assembly. Additionally, a native promoter-driven *trxA* gene was incorporated to this plasmid for complementation. For the expression of Cas12 hacker RNP in *E*. *coli*, 6×His-SUMO-tagged Cas12 hacker and a T7-driven sgRNA_v1 expression cassette were inserted into the pET28a-based vector. Expression plasmids for mutant and truncated versions of Cas12 hacker were constructed using either Gibson assembly or circular polymerase extension cloning (CPEC)^77^. For the protein expression of TrxA in *E*. *coli*, an N-terminal 6×His-tagged *trxA* coding sequence were cloned into a pET28a-based vector. For the *in vitro* dsDNA cleavage assay, the plasmid for DNA substrate were constructed by incorporating protospacer and different PAM sequences into pUC19-based vector using Gibson assembly and CPEC. For the pull-down assay, a T7 lac-driven Cas12 hacker nuclease and its non-coding region was cloned into the TrxA protein expression plasmid. Mutations in Cas12 hacker or *trxA* gene were introduced via Gibson assembly. For the pull-down assay of His-TrxC and untagged Cas12 hacker, the plasmid was constructed by replacing the *trxA* gene with *trxC* gene. Detailed information of the plasmids is listed in Supplementary Table 5.

### Preparation of electrocompetent cells

For plasmid-free electrocompetent cells of *E*. *coli* BL21(DE3), *K. pneumoniae* 1.6366, and *A*. *baumannii* 17978, 1 mL of the overnight culture was diluted into 100 mL of fresh LB broth and incubated at 37°C until the culture reached an optical density at 600 nm (OD_600_) of 0.5. Cells were harvested by centrifugation at 4°C and the supernatant was discarded. The cell pellet was gently resuspended in 20 mL of sterile ice-cold 10% glycerol. This process of centrifugation and resuspension was repeated twice. Finally, the cell pellet was resuspended in 1 mL of sterile ice-cold 10% glycerol and dispensed into 50 μL aliquots for electroporation.

For PAM depletion assay, the plasmid harboring CRISPR-Cas12 hacker locus was transformed into *E*. *coli* DH5α cells. Then, 1 mL of overnight culture was diluted into 100 mL of LB broth containing 50 μg/mL apramycin and incubated at 37°C with shaking until the OD_600_ reached 0.5. Subsequent procedures were consistent with those described above for plasmid-free electrocompetent cells.

For bacterial genome interference assay, the Cas12 hacker-harboring plasmid was transformed into plasmid-free electrocompetent cells *E*. *coli* BL21(DE3), *K. pneumoniae* 1.6366, and *A*. *baumannii* 17978, respectively. The subsequent procedures for producing electrocompetent cells of *E*. *coli* and *K. pneumoniae* were carried out in a manner as described above for PAM depletion assay. For the preparation of *A*. *baumannii* electrocompetent cells, 1 mM Isopropyl-β-D-thiogalactopyranoside (IPTG) was used to induce the expression of Cas12 hacker nuclease when the OD_600_ reached approximately 0.2, and other procedures were consistent with those described above for PAM depletion assay.

### PAM depletion assay and analysis

100 ng of the PAM library plasmids were electroporated into the *E. coli* DH5α electrocompetent cells expressing the CRISPR-Cas12 hacker system. The transformants were plated on the LB agar containing 50 μg/mL apramycin and 50 μg/mL carbenicillin. Colonies were harvested for plasmid extraction by SPARKeasy Superpure Mini Plasmid Kit (Shandong Sparkjade Biotechnology Co., Ltd). The amplicon sequencing library was constructed by a two-step amplification method. The PCR product of PAM randomized region was purified using TIANquick Midi Purification Kit (Tiangen Biotech, Beijing, China). Then, the amplicon sequencing library was prepared using NEBNext® Ultra™ II DNA Library Prep Kit (Vazyme Biotech) and sequenced on an Illumina Novaseq 6000 platform at HaploX Genomics Center (Jiangxi, China).

For data analysis, the number of each PAM sequence in the randomized regions was counted. Subsequently, the proportion of each PAM in the experimental group (targeted by Cas12 hacker) and the control group (non-targeted) was calculated. The log ratio of PAM depletion values between the experimental and control groups was then determined. By comparing these ratios, the PAM specificity of Cas12 hacker was characterized. PAM sequences were visualized using WebLogo and heatmap.

### Bacterial genome interference assay

For the bacterial genome interference assay depicted in Fig. 5b and Supplementary Fig. 14b,c, 100 ng of the plasmids containing sgRNA_v2 cassette (with or without target) were electroporated into 50 μL electrocompetent cells of different *E*. *coli* BL21(DE3) strains, which already contained Cas12 hacker-harboring plasmids. The electroporation was conducted using a 2 mm cuvette with parameters set as 2.5 kV, 200 Ω, and 25 μF (Bio-Rad). After electroporation, the cells were recovered in 1 mL LB medium for 1.5 hours at 37 ℃. Subsequently, the cells were centrifuged and resuspended with 100 μL LB, and then the resuspended cells were subjected to consecutive 10-fold dilutions. The diluted bacterial cells were spotted on the LB agar plates supplemented with 50 μg/mL apramycin and 50 μg/mL kanamycin and cultured at 37°C overnight. The genome interference assays in *K*. *pneumoniae* 1.6366 and *A*. *baumannii* 17978 strains were performed following the same procedure.

For the bacterial genome interference assay depicted in Fig. 5c, plasmids containing the sgRNA_v2 expression cassette or the sgRNA_v2 and *trxA* gene expression cassettes were electroporated into 50 μL *E*. *coli* BL21(DE3) *trxA*- deletion electrocompetent cells that harbored a plasmid with Cas12 hacker. The subsequent operations followed those described above.

For the bacterial genome interference assay depicted in Supplementary Fig. 13d, 100 ng of the plasmids containing sgRNA_v2 and wild-type or mutant *trxA* expression cassette were electroporated into 50 μL *E*. *coli* BL21(DE3) *trxA*- deletion electrocompetent cells that harbored a plasmid with Cas12 hacker. The subsequent operations followed those described above.

For the bacterial genome interference assay depicted in Supplementary Fig. 13f, 100 ng of the sgRNA_v2-harboring plasmids and 150 ng of the wild-type or mutant Cas12 hacker-harboring plasmids were co-electroporated into 50 μL of wild-type *E*. *coli* BL21(DE3) cells. The electroporation operation followed those described above. After recovery, the cells were centrifuged and resuspended with 50 μL of LB, and the resuspended cells were then subjected to consecutive 5-fold dilutions. The diluted cells were spotted on the LB agar plates supplemented with 50 μg/mL apramycin and 50 μg/mL kanamycin, then incubated at 37°C overnight.

### Construction of mutant strains

*E*. *coli* BL21(DE3) *trxA*-deletion mutant, *E*. *coli* BL21(DE3) *trxC*-deletion mutant, and *K*. *pneumoniae* 1.6366 *trxA*- deletion mutant were produced by the pCasKP-pSGKP two-plasmid system, following previously described procedures^75^. The *A*. *baumannii* 17978 *trxA*-deletion mutant was generated by the pCasAb/pSGAb two-plasmid system following the procedures described previously^76^. The spacers and the repair templates used for gene deletion were listed in Supplementary Table 5.

### Phage plaque assays

The p15A-based plasmids expressing Cas12 hacker and sgRNA_v2 targeting T4 or T7 phage genome were transformed into *E*. *coli* BL21(DE3) competent cells. The transformed cells were shaken at 37°C overnight in LB medium supplemented with 50 μg/mL apramycin. Next day, 0.3 mL overnight bacterial culture was mixed with 3 mL soft LB agar with 50 μg/mL apramycin and poured on 90-mm^2^ LB agar plate containing 50 μg/mL apramycin. The phage was subjected to consecutive 10-fold dilutions in LB medium, and 3 μL of the diluted phage was spotted on the solidified agar. The plates were then incubated at 37°C.

For assessing the role of *trxA* in the CRISPR-Cas12 hacker system against phage infection, the p15A-based plasmids expressing Cas12 hacker and sgRNA_v2, or the modified p15A plasmids expressing Cas12 hacker, *trxA*, and sgRNA-v2 were transformed into *E*. *coli* BL21(DE3) *trxA*-deletion competent cells. The subsequent experimental process was identical to that described above. All experiments were performed in triplicate. Target sequences can be found in Supplementary Table 5.

### Pull-down assay

For the pull-down assay in Fig. 1i, the plasmid encoding the CRISPR-Cas12 hacker locus and His-TxrA (or His-TrxC) was transformed into BL21(DE3) cells deficient in *trxA* (or *trxC*). The seed cultures were then inoculated into 15 mL LB broth with 50 μg/mL kanamycin and incubated at 37°C until the OD_600_ reached 0.6. Protein expression was induced by adding 0.3 mM IPTG and incubating the culture at 16°C overnight. Cells were collected and resuspended in buffer F (20 mM Tris-HCl, pH 7.5, 300 mM NaCl, 10 mM imidazole, and 5% glycerol). Then, the cells were lysed by sonication, and the lysates were cleared by centrifugation. The resulting supernatants were incubated with 50 μL of Ni Smart Magarose Beads (Smart-Lifesciences) at 4 °C for 2 h. The beads were then washed three times with 1 mL of buffer F to remove unbound proteins. Finally, the proteins were eluted by washing the beads with 50 μL of buffer G (20 mM Tris-HCl, pH 6.8, 300 mM NaCl, 300 mM imidazole, and 5% glycerol). The eluates were further analyzed by reducing SDS-PAGE using SurePAGE™ precast gel (GenScript) and stained with Coomassie brilliant blue (CBB).

For the pull-down assay in Fig. 4d,e, Cas12 hacker or TrxA mutants were introduced into the plasmid expressing the CRISPR-Cas12 hacker locus and His-TxrA. These plasmids were then transformed into BL21(DE3) cells deficient in *trxA*. The subsequent steps were the same as those described for Fig. 1i.

For pull-down assay in Fig. 4f and Supplementary Fig. 13g, cysteine mutants of Cas12 hacker and TrxA (TrxA-C33S, TrxA-C36S, Cas12 hacker-C127S) were introduced into the plasmid expressing the CRISPR-Cas12 hacker locus and His- TxrA. Then, these plasmids were transformed into BL21(DE3) cells deficient in *trxA*. The procedures of protein expression and elution were the same as those described for Fig. 1i. Lastly, the eluates were divided into two parts and respectively subjected to reducing and non-reducing SDS-PAGE analysis, followed by CBB staining.

### Protein expression and purification

For expressing Cas12 hacker-sgRNA_v1 RNP complex, the pET28a-based RNP expression plasmid was transformed into *E*. *coli* BL21(DE3), and the cells were plated on the LB plate containing 50 μg/mL kanamycin. 10 mL of overnight culture was inoculated into 1 L fresh LB broth with 50 μg/mL kanamycin and shaken at 37°C until the OD_600_ reached 0.6. Protein expression was induced by adding IPTG to a final concentration of 0.3 mM. Subsequently, the cells were cultured at 16°C overnight and collected by centrifugation.

The cell pellets were resuspended in buffer A (20 mM Tris-HCl, pH 7.5, 300 mM NaCl, 10 mM imidazole, 5% glycerol, 1 mM PMSF, and 0.5 mM TCEP), then lysed by sonication. After centrifugation, the supernatant was loaded onto a 5-mL HisTrap HP column (GE healthcare/Cytiva). The column was washed with 20 mL of buffer B (20 mM Tris-HCl, pH 7.5, 300 mM NaCl, 62.5 mM imidazole, 5% glycerol, 0.2 mM PMSF, and 0.5 mM TCEP) to remove unbound proteins, and then eluted with buffer C (20 mM Tris-HCl, pH 6.8, 300 mM NaCl, 500 mM imidazole, 5% glycerol, 0.2 mM PMSF, and 0.5 mM TCEP). The eluates were further purified by using a HiLoad 16/600 Superdex 200 pg column (GE healthcare/Cytiva) with buffer D (20 mM Tris-HCl, pH 7.5, 300 mM NaCl, and 0.5 mM TCEP). The final product was concentrated to about approximately 10 μM, which is quantified by *Easy* Protein Quantitative Kit (Bradford) (TransGen Biotec) and subsequently stored at -80°C after rapid freezing in liquid nitrogen. All purification steps were carried out at 4°C. The expression and purification processes of the mutated and truncated Cas12 hacker RNP complexes were the same as that of the Cas12 hacker-sgRNA_v1 RNP complex.

For the purification of Cas12 hacker-sgRNA_v1 RNP complex without TrxA or TrxC, the pET28a-based RNP expression plasmid was transformed into the competent cells of *E*. *coli* BL21(DE3) *trxA*-deletion or *trxC*-deletion strain. For the purification of His-TrxA protein, the pET28a-based TrxA expression plasmid was transformed into *E*. *coli* BL21(DE3). The subsequent purification steps were conducted as described above.

### *In vitro* dsDNA cleavage assay

For the cleavage of 56-bp dsDNA labeled with 5’ FAM and 3’ Texas Red, a 10 μL reaction mixture was prepared with 20 nM DNA substrate, 500 nM Cas12 hacker-sgRNA_v1 RNP complex, 10 mM Tris-HCl, pH 7.5, 10 mM MgCl2, and 50 mM NaCl. The mixture was incubated at 37°C and subsequently quenched with 10 μL formamide stop buffer (90% formamide, 50 mM EDTA, 0.075% bromophenol blue, and 0.075% xylene cyanol FF). After a 10-minute incubation at 95°C, the cleavage products were analyzed by electrophoresis using 20% TBE-Urea-PAGE gel.

For the cleavage of 2.7-kb dsDNA, the linear DNA substrate was prepared by PCR amplification using 2× Hieff Canace® AdvanceFast PCR Master Mix (With Dye) (Yeasen) and then purified with SPARKeasy Gel/PCR Purification Kit (Shandong Sparkjade Biotechnology Co., Ltd). A 10 μL reaction mixture was prepared, containing 5 nM linear DNA substrate, 300 nM Cas12 hacker-sgRNA_v1 RNP complex, 10 mM Tris-HCl, pH 7.5, 10 mM MgCl_2_, and 50 mM NaCl. The mixture was incubated at 37°C and terminated by adding 6× Gel Loading Dye (NEB) containing 25 mM EDTA and 0.5 μL of Proteinase K (20 mg/ml). After a 10-minute incubation at 55°C, the cleavage products were analyzed by 1% agarose gel electrophoresis.

### Small RNA-seq and analysis

The p15A-based plasmid encoding the CRISPR-Cas12 hacker locus was transformed into *E*. *coli* DH5α strain. Total RNA was isolated using the TaKaRa MiniBEST Universal RNA Extraction Kit (TaKaRa). To eliminate DNA contamination, 15 μg RNA was treated with 2 units of DNase I (NEB) at 37°C for 30 min, followed by phenol/chloroform extraction. The RNA was then subjected to 2’-3’-dephosphorylation by incubating with 20 units of T4 PNK (NEB) at 37°C for 6 hours. Subsequently, 1 mM ATP was added, and the mixture was incubated at 37°C for 1 h to achieve 5’-phosporylation. The phenol/chloroform purified RNA was treated with 5 units of RppH (NEB) at 37°C for 1 h to hydrolyze the 5’ pyrophosphate. Following another phenol/chloroform extraction, the pre-treated RNA was qualified by agarose gel electrophoresis and used to construct the small RNA library using VAHTS Small RNA Library Prep Kit for Illumina (Vazyme Biotech) according to the manufacturer’s instruction. Then, the RNA library was sequenced on the Illumina HiSeq platform (PE150) at HaploX Genomics Center (Jiangxi, China). The sequencing reads were mapped to the reference sequence using HISAT2^78^. The mapped reads were analyzed using SAMtools^79^. For the RNA extracted from the Cas12 hacker-sgRNA_v3 RNP complex, the similar methods for sample preparation and results analysis were applied.

### Mass spectrometric analysis

Mass spectrometric analysis for the sample of the purified Cas12 hacker-sgRNA_v1 RNP was performed by Mass Spectrometry System at the National Facility for Protein Science in Shanghai (NFPS). The protein sample was first digested to peptide mixture, which was then loaded onto an Easy-nLC1200 system equipped with a home-made reverse phase C18 column. A 120-minute gradient from 2% to 100% of buffer B (buffer A: 0.1% formic acid in water; buffer B: 0.1% formic acid in 80% Acetonitrile) was applied at a flow rate of 300 nL/min. The eluted peptides were ionized and directly introduced into a Q-Exactive mass spectrometer (Thermo Scientific, San Jose, CA) via a nano-spray source using a distal 2.5-kV spray voltage. MS scan functions and LC solvent gradients were controlled by the Xcalibur data system (Thermo Scientific). The acquired MS/MS data were analyzed against a homemade database (including all target proteins) using PEAKS Studio 8.5. With the raw data, the way we screened for candidate proteins was that the molecular weight size of target proteins was between 10-20 kDa, and the proportion was also relatively high. The raw data are shown in Supplementary Table 2.

### Cryo-EM sample preparation and data collection

For cryo-EM sample preparation, the Cas12 hacker-sgRNA-target DNA complex was assembled by mixing the purified Cas12 hacker-sgRNA_v1 RNP with the target DNA at a molar ratio of 1:1.3 in buffer E (20 mM Tris-HCl, pH 7.5, 300 mM NaCl, 5 mM MgCl2, and 0.5 mM TCEP). Two types of DNA substrates were used: one consisting of a 29-nt target strand paired with an 11-nt non-target strand, and the other being a 33-bp double-stranded DNA. The ternary complex was first incubated at 25°C for 20 minutes, followed by a 10-minute incubation at 37°C. Subsequently, the complex was further purified using a Superose 6 increase 10/300 GL column (GE healthcare/Cytiva) which was pre-equilibrated with buffer E. The ternary complex with the staggered DNA was concentrated to approximately 3 mg/mL, while the complex with the blunt DNA was concentrated to about 3.6 mg/mL.

The film grids (Au 300 mesh, R1.2/1.3, ANTcryo, Guangzhou) were first glow-discharged for 30 s in a Solarus (950) plasma cleaning system (Gatan, USA) with H2/O2. Subsequently, 4 μL of the complex was applied onto the glow-discharged grid. The grid was then blotted for 3 s at 4 °C under 100% humidity with a blot force of -1. Following this, the grid was flash-frozen in liquid ethane using a Vitrobot Mark IV (Thermo Fisher Scientific).

Cryo-EM data were collected at a 300 KV Titan Krios electron microscope (Thermo Fisher Scientific, USA) equipped with a K3 detector (Gatan, USA). The defocus range was set between -1.2 μm and -1.8 μm, and a total dose of 60 electrons per Å^2^. 3,976 micrographs for the Cas12 hacker-sgRNA-target DNA complex with staggered DNA and 8,004 micrographs for the complex with blunt DNA were collected using SerialEM^80^ at a magnification of 105000×, corresponding to a calibrated pixel size of 0.832 Å. A full description of the cryo-EM data collection parameters can be found in Supplementary Table 4.

### Cryo-EM data processing

The movies were subjected to beam-induced motion correction using the MotionCor2 algorithm implemented in RELION^81^. The following data processing was performed using cryoSPARC^82^. The contrast transfer function (CTF) parameters were estimated by patch-based CTF estimation. Particles were automatically picked by blob picker, and extracted for 2D classifications. The particles were subsequently subjected to ab-initio reconstruction and curated by heterogenous refinement. The selected particles were subjected to non-uniform refinement^83^. The local resolution estimation was also performed in cryoSPARC. DeepEMhancer^84^ was used for post-processing to generated the sharpen map.

### Model building and refinement

For the quaternary Cas12 hacker-TrxA-sgRNA-target DNA (29-nt TS and 11-nt NTS) complex involving the partial TB domain, the protein and nucleic acids models were manually built by COOT^85^, guided by the predicted models of Cas12 hacker and TrxA generated by AlphaFold2. The model was refined using the phenix.real_space_refine tool in PHENIX^86^, with secondary structure restraints. For the quaternary Cas12 hacker-TrxA-sgRNA-target DNA (33-bp dsDNA) complex involving the intact TB domain, the model was built using the model with partial TB domain as a reference. The cryo-EM map and atomic models were visualized with USCF ChimeraX^87^.

## Data availability

The structures of Cas12 hacker-TrxA-sgRNA-target DNA have been deposited in the Protein Data Bank under the accession numbers 9JFS and 9JG3, and in the EMDB under the accession numbers EMD-61438 and EMD-61449. The raw data of PPI screening and MS are provided in Supplementary Tables 1 and 2, respectively. The next-generation sequencing data containing PAM depletion and RNA-seq raw reads are available on the Sequence Read Archive under BioProject PRJNA1159697 and PRJNA1159999.

## Acknowledgments

We thank the support from the Bio-EM facility at ShanghaiTech University for cryo-EM data collections, and we are grateful to Li Wang and Dandan Liu for their help with cryo-EM technical support. We thank Chen Su of the Mass Spectrometry System at the National Facility for Protein Science in Shanghai (NFPS), Shanghai Advanced Research Institute, Chinese Academy of Science, China for MS sample preparation, data collection, and data analysis. The work was supported by HPC Platform of ShanghaiTech University. ChatGPT-4 was used for language editing. This work was supported by grants 2022YFC3403400 and 2023YFC3400200 from the National Key R&D Program of China, 22ZR1480100, and 23HC1400800 from the Shanghai Committee of Science and Technology, China, 22277078 and 22077083 from the National Natural Science Foundation of China, 2023M742367 and 2024M752064 from China Postdoctoral Science Foundation, and GZC20231672 from Postdoctoral Fellowship Program of CPSF.

## Author Contributions

Q.J. directed the project. Q.J., Z.W., and Y.W. conceived the project. Z.W. established the CRISPR-Cas12 systems mining pipeline and conducted the PPIs and bioinformatics analyses. Z.W. and Y.W. constructed the plasmids, and performed the PAM-depletion and small RNA-seq assays. Y.W. performed the phage plaque, pull-down, and *in vitro* DNA cleavage assays. Y.W., Z.W., H.G., and J.D. performed the genome interference assay. Y.W., Z.W., H.G., and Ya.W. performed the protein purification. Z.W., Y.W., N.T., H.G., and J.D. prepared the cryo-EM samples, and Z.W. determined the structures. Q.J., Z.W., and Y.W analyzed and discussed the experimental data. Z.W. and Y.W prepared the figures. Q.J., Z.W., and Y.W wrote the paper.

