## Supplementary information for "AI-driven discovery of host thioredoxin as a CRISPR enhancer of phage-encoded miniature Cas12 hacker nuclease"

**This file includes:**

**Supplementary Table 1. PPI screening results of candidate cas12 proteins and *Escherichia coli* proteome**
**Supplementary Table 2. Mass spectrometry results of the co-purified proteins**
**Supplementary Table 3. Taxonomic summary of CRISPR-Cas12 hacker loci using Virsorter 2 and geNomad**
**Supplementary Table 4. Statistics for cryo-EM analysis and model refinement**
**Supplementary Table 5. Plasmids, targets, nucleotides, and proteins information**
**Supplementary Fig. 1 Protein-protein interaction (PPI) screening results of candidate Cas12 (CC12) proteins containing potential intrinsically disordered regions (IDRs)**
**Supplementary Fig. 2 Bioinformatics analysis of the CRISPR-Cas12 hacker family**
**Supplementary Fig. 3 PAM preferences and biochemical properties of Cas12 hacker**
**Supplementary Fig. 4 Cryo-EM data analysis of the Cas12 hacker-sgRNA-target DNA (29-nt TS and 11-nt NTS) complex**
**Supplementary Fig. 5 Complex structures of different proteins and thioredoxin**
**Supplementary Fig. 6 Comparison of different domains in TnpB, Cas12k, and Cas12 hacker**
**Supplementary Fig. 7 Structural comparison of TnpB and Cas12 proteins in complex with corresponding sgRNA/ωRNA and target DNA**
**Supplementary Fig. 8 Structural comparison of WED, REC, and RuvC domains for TnpB, Cas12 hacker, and other Cas12 proteins**
**Supplementary Fig. 9 Comparative analysis of sgRNA-target DNA architecture**
**Supplementary Fig. 10 Schematic for guide RNA and target DNA recognition**
**Supplementary Fig. 11 Recognition of sgRNA and target DNA by Cas12 hacker**
**Supplementary Fig. 12 Comparison of the PI domain of Cas12 hacker and Cas12k**
**Supplementary Fig. 13 Characterization of the interaction between TrxA and Cas12 hacker**
**Supplementary Fig. 14 The essential role of TrxA in modulating Cas12 hacker's activity**
**Supplementary Fig. 15 Cryo-EM data analysis of the Cas12 hacker-TrxA-sgRNA-target DNA (33-bp dsDNA) complex**
**Supplementary Fig. 16 The mechanism of Cas12 hacker's TB domain**

**Supplementary Table 4. Statistics for cryo-EM analysis and model refinement**

| **Sample** | Cas12 hacker-TrxA-sgRNA-target DNA (29-nt TS and 11-nt NTS) | Cas12 hacker-TrxA-sgRNA-target DNA (33-bp dsDNA) |
|---|---|---|
| EMDB ID | EMD-61438 | EMD-61449 |
| PDB ID | 9JFS | 9JG3 |
| **Data collection** | | |
| Microscope | Titan Krios | Titan Krios |
| Voltage (kV) | 300 | 300 |
| Detector | Gatan K3 | Gatan K3 |
| Magnification | 105,000 | 105,000 |
| Voltage (kV) | 300 | 300 |
| Pixel size (Å) | 0.832 | 0.832 |
| Electron exposure (e-/Å2) | 60 | 60 |
| Defocus range (μm) | -1.2 to -1.8 | -1.2 to -1.8 |
| Number of movies | 3,976 | 8,004 |
| **Reconstruction** | | |
| Initial particle images | 3,094,626 | 3,039,656 |
| Final particle images | 508,876 | 66,342 |
| Map resolution (Å) | 2.67 | 3.20 |
| FSC threshold | 0.143 | 0.143 |
| Symmetry | *C1* | *C1* |
| **Model building and refinement** | | |
| Model composition | | |
| Non-hydrogen atoms | 8796 | 9188 |
| Protein atoms | 634 | 684 |
| Nucleic acid atoms | 175 | 174 |
| R.m.s deviations | | |
| Bond length (Å) | 0.005 | 0.005 |
| Bond angle (˚) | 0.532 | 0.615 |
| Validation | | |
| Clash score | 3.56 | 7.94 |
| Rotamer outliers (%) | 0.53 | 0.00 |
| Ramachandran plot | | |
| Favored (%) | 97.76 | 98.08 |
| Allowed (%) | 2.24 | 1.92 |
| Outlier (%) | 0.00 | 0.00 |

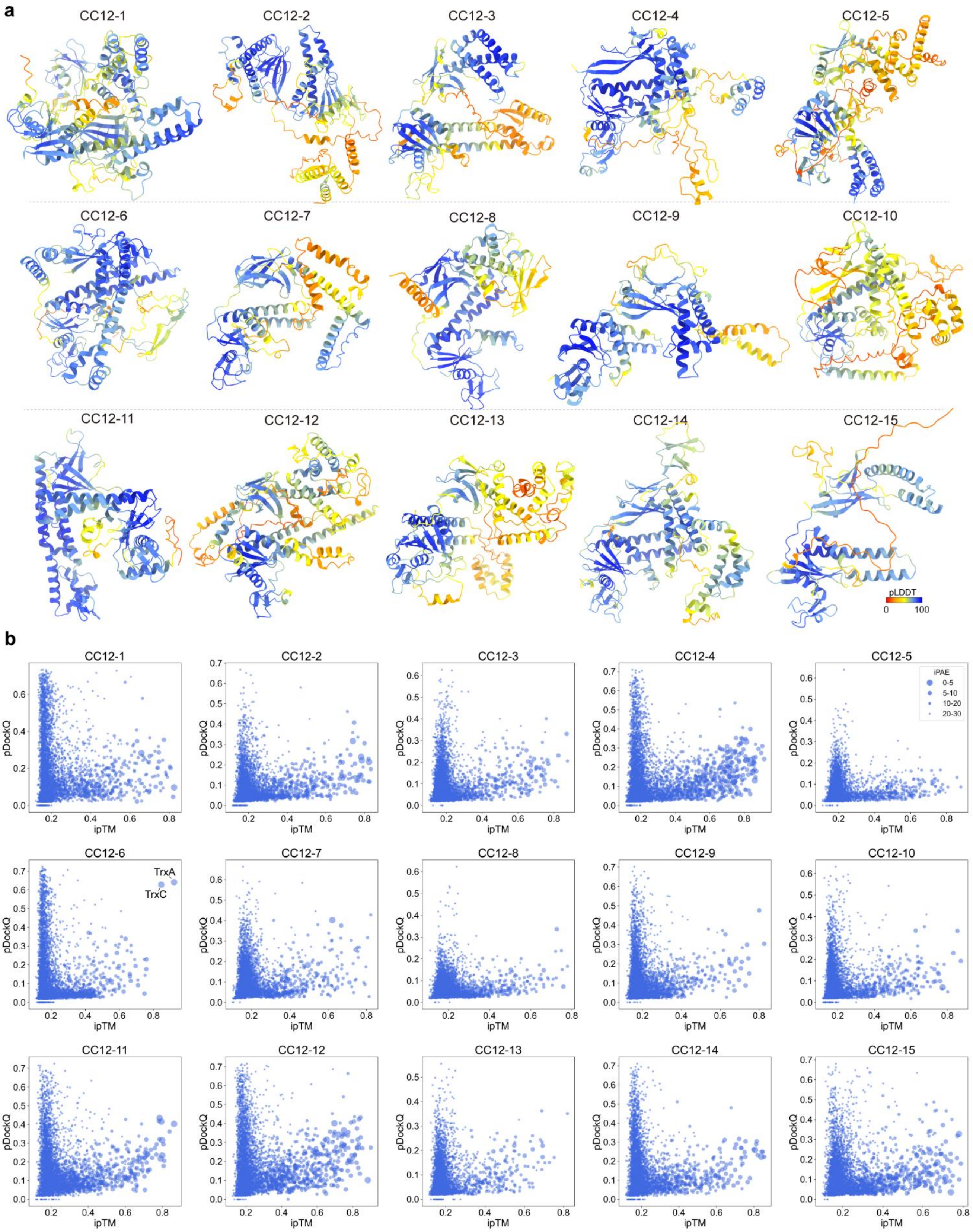


**Supplementary Fig. 1 Protein-protein interaction (PPI) screening results of candidate Cas12 (CC12) proteins containing potential intrinsically disordered regions (IDRs)**

**a,** Top-ranking models of 15 candidate Cas12 proteins containing potential IDRs. The structural models were predicted by AlphaFold2 and colored by pLDDT scores.

**b,** The bubble plots visualized the PPIs prediction results of 15 CC12 proteins and the *E. coli* proteome. The scoring metrics for PPIs screening include ipTM, pDockQ, and iPAE. The X-axis, the Y-axis, and varing dot size represent ipTM, pDockQ, and iPAE, respectively.

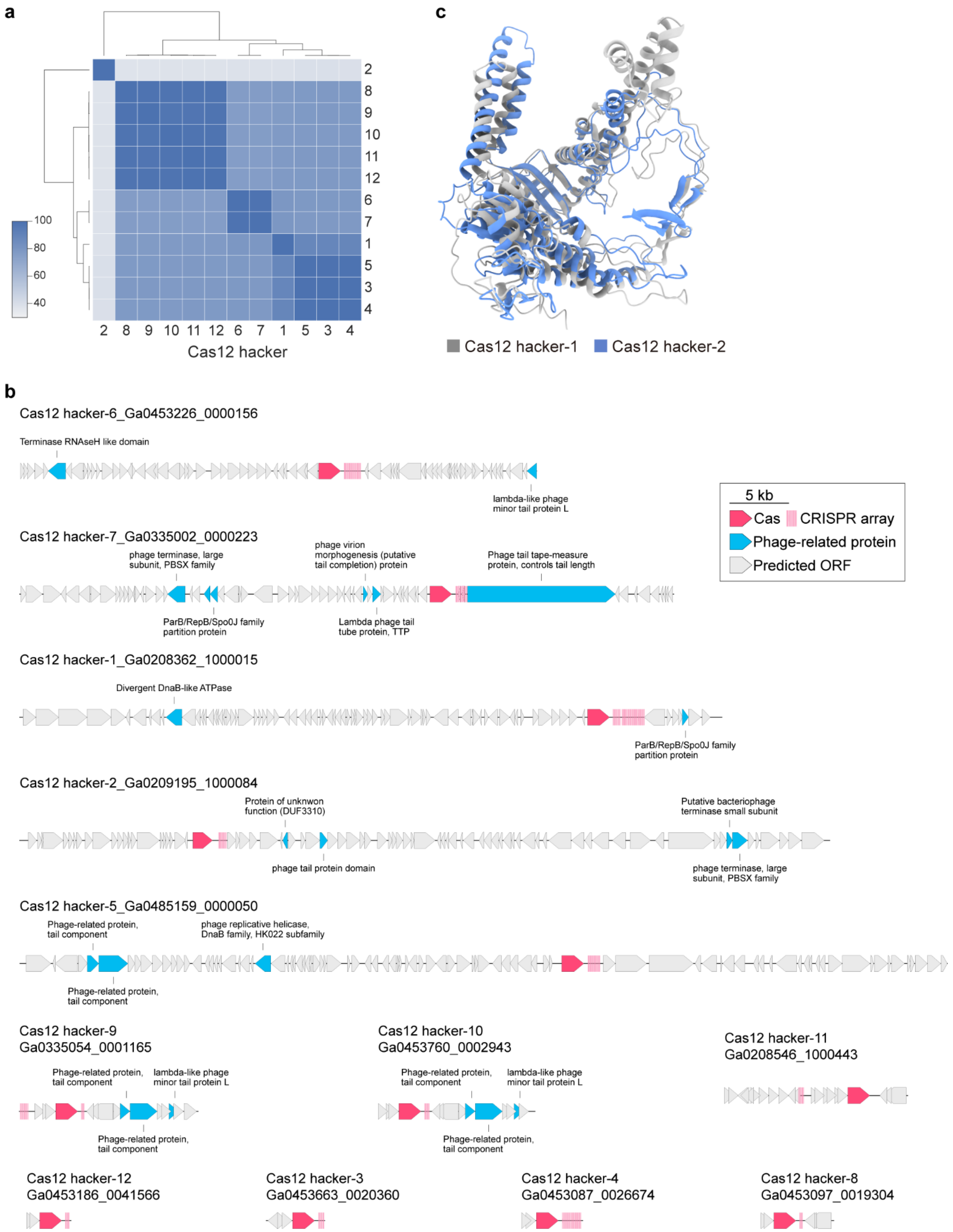

a
2 8 9 10 11 12 6 7 1 5 3 4
Cas12 hacker
100
80
60
40
c
Cas12 hacker-1
Cas12 hacker-2
b
Cas12 hacker-6_Ga0453226_0000156
Terminase RNAseH like domain
lambda-like phage minor tail protein L
Cas12 hacker-7_Ga0335002_0000223
phage terminase, large subunit, PBSX family
phage virion morphogenesis (putative tail completion) protein
Phage tail tape-measure protein, controls tail length
ParB/RepB/Spo0J family partition protein
Lambda phage tail tube protein, TTP
5 kb
Cas
CRISPR array
Phage-related protein
Predicted ORF
Cas12 hacker-1_Ga0208362_1000015
Divergent DnaB-like ATPase
ParB/RepB/Spo0J family partition protein
Cas12 hacker-2_Ga0209195_1000084
Protein of unknwon function (DUF3310)
Putative bacteriophage terminase small subunit
phage tail protein domain
phage terminase, large subunit, PBSX family
Cas12 hacker-5_Ga0485159_0000050
Phage-related protein, tail component
phage replicative helicase, DnaB family, HK022 subfamily
Phage-related protein, tail component
Cas12 hacker-9
Ga0335054_0001165
Phage-related protein, tail component
lambda-like phage minor tail protein L
Phage-related protein, tail component
Cas12 hacker-10
Ga0453760_0002943
Phage-related protein, tail component
lambda-like phage minor tail protein L
Phage-related protein, tail component
Cas12 hacker-11
Ga0208546_1000443
Cas12 hacker-12
Ga0453186_0041566
Cas12 hacker-3
Ga0453663_0020360
Cas12 hacker-4
Ga0453087_0026674
Cas12 hacker-8
Ga0453097_0019304

**Supplementary Fig. 2 Bioinformatics analysis of the CRISPR-Cas12 hacker family**

**a**, Sequence identity analysis of the Cas12 hacker proteins. Except for Cas12 hacker-2, the other proteins show high sequence identity.

**b**, Comprehensive architecture of metagenomic assemblies encompassing CRISPR-Cas12 hacker loci. The annotations for hallmark genes are generated by Genomad. Cas and CRISPR array, phage-related protein, and predicted ORF are indicated in red, blue, and grey, respectively.

**c**, Structural alignment of Cas12 hacker-1 and Cas12 hacker-2. The structures of Cas12 hacker-1 and Cas12 hacker-2 are predicted by AlphaFold2, and coloured in grey and blue, respectively.

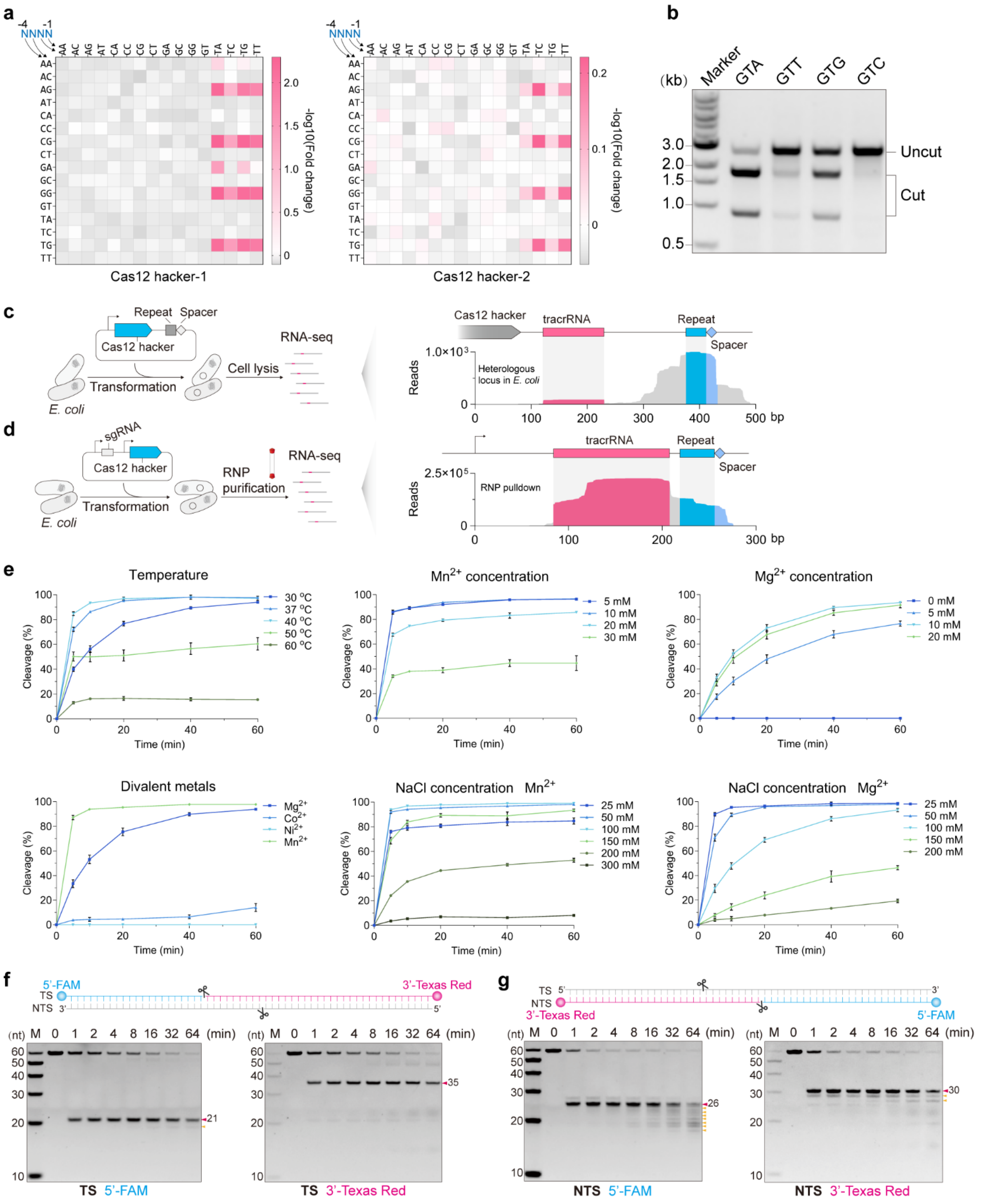


**Supplementary Fig. 3 PAM preferences and biochemical properties of Cas12 hacker**

**a**, 4N Heatmap of PAM sequence for Cas12 hacker-1 (left) and Cas12 hacker-2 (right). Cas12 hacker-1 shows a higher PAM depletion efficiency than Cas12 hacker-2.

**b**, *In vitro* dsDNA cleavage with four different PAMs by Cas12 hacker-1. The order of the preferred PAM sequence is

GTA > GTG > GTT > GTC.

**c**,**d,** Schematics and results of small RNA sequencing for the CRISPR-Cas12 hacker from *E. coli* heterologous expression (**c**) or RNP pulldown (**d**).

**e**, Detailed cleavage kinetics of Cas12 hacker under various conditions. This panel presents the cleavage efficiency of Cas12 hacker across different experimental variables, including temperature, the presence of divalent metal ions, concentrations of $Mg^{2+}$ and $Mn^{2+}$, and NaCl concentration, offering insights into the optimal biochemical conditions for its activity.

**f,g,** TBE-Urea-PAGE assays demonstrates the precise cleavage patterns of FAM/Texas red-labelled TS DNA (**f**) and NTS DNA (**g**) by Cas12 hacker.

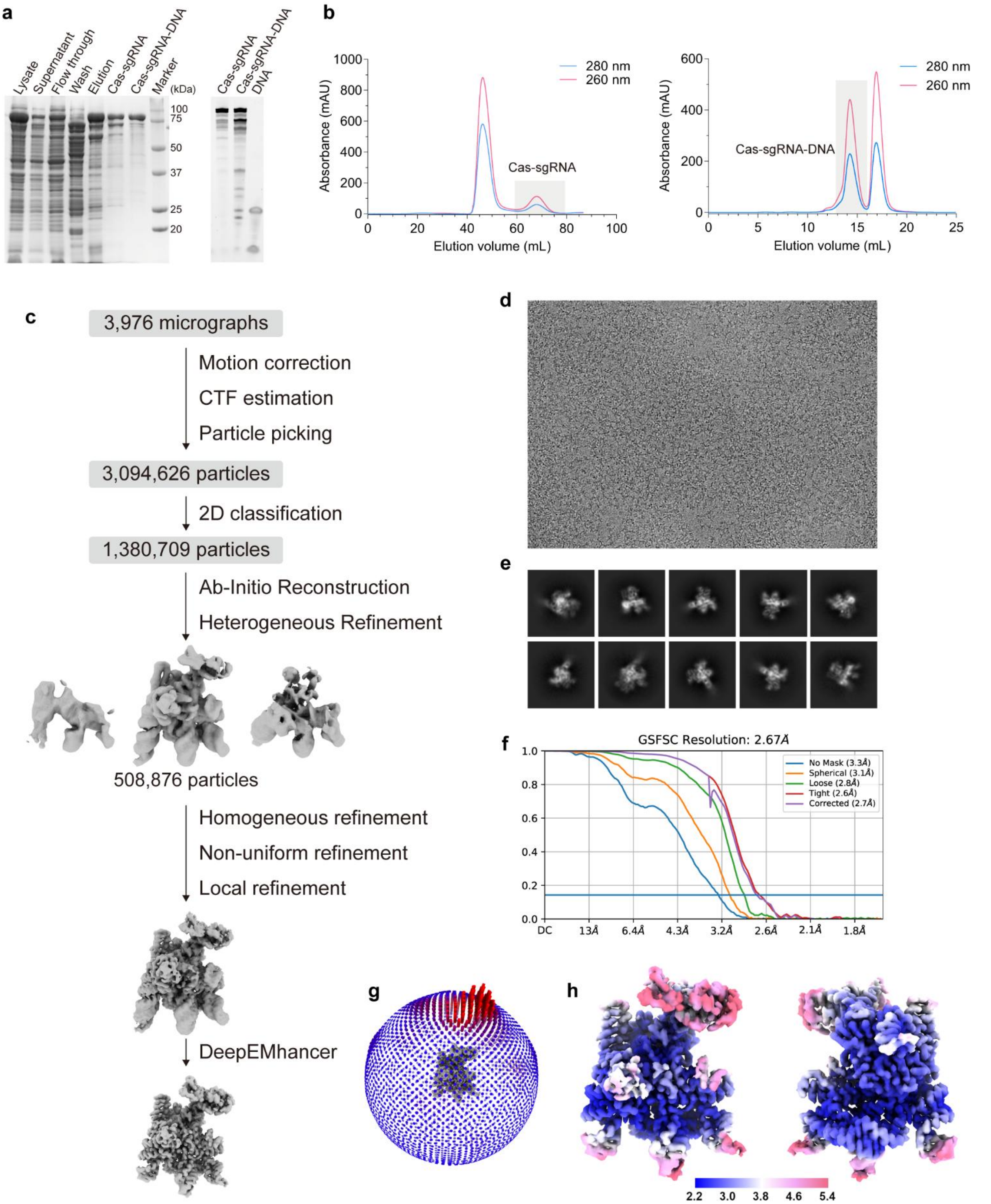


**Supplementary Fig. 4 Cryo-EM data analysis of the Cas12 hacker-sgRNA-target DNA (29-nt TS and 11-nt NTS) complex**

**a**, SDS-PAGE (stained with CBB, left) and Urea-PAGE (right) analysis of the Cas12 hacker-sgRNA-target DNA (29-nt TS and 11-nt NTS) complex.

**b**, Size-exclusion chromatography (SEC) profile of Cas12 hacker-sgRNA RNP (left) and the Cas12 hacker-sgRNA-target DNA (29-nt TS and 11-nt NTS) complex (right).

**c**, Cryo-EM data processing workflow for the Cas12 hacker-sgRNA-target DNA (29-nt TS and 11-nt NTS) complex.

**d**, Representative cryo-EM micrograph from a total of 3976 micrographs.

**e**, Representative 2D class averages.

**f**, The half-map Fourier shell correlation (FSC) curves.

**g**, Euler angle distribution of particles.

**h**, Cryo-EM density map coloured by local resolution.

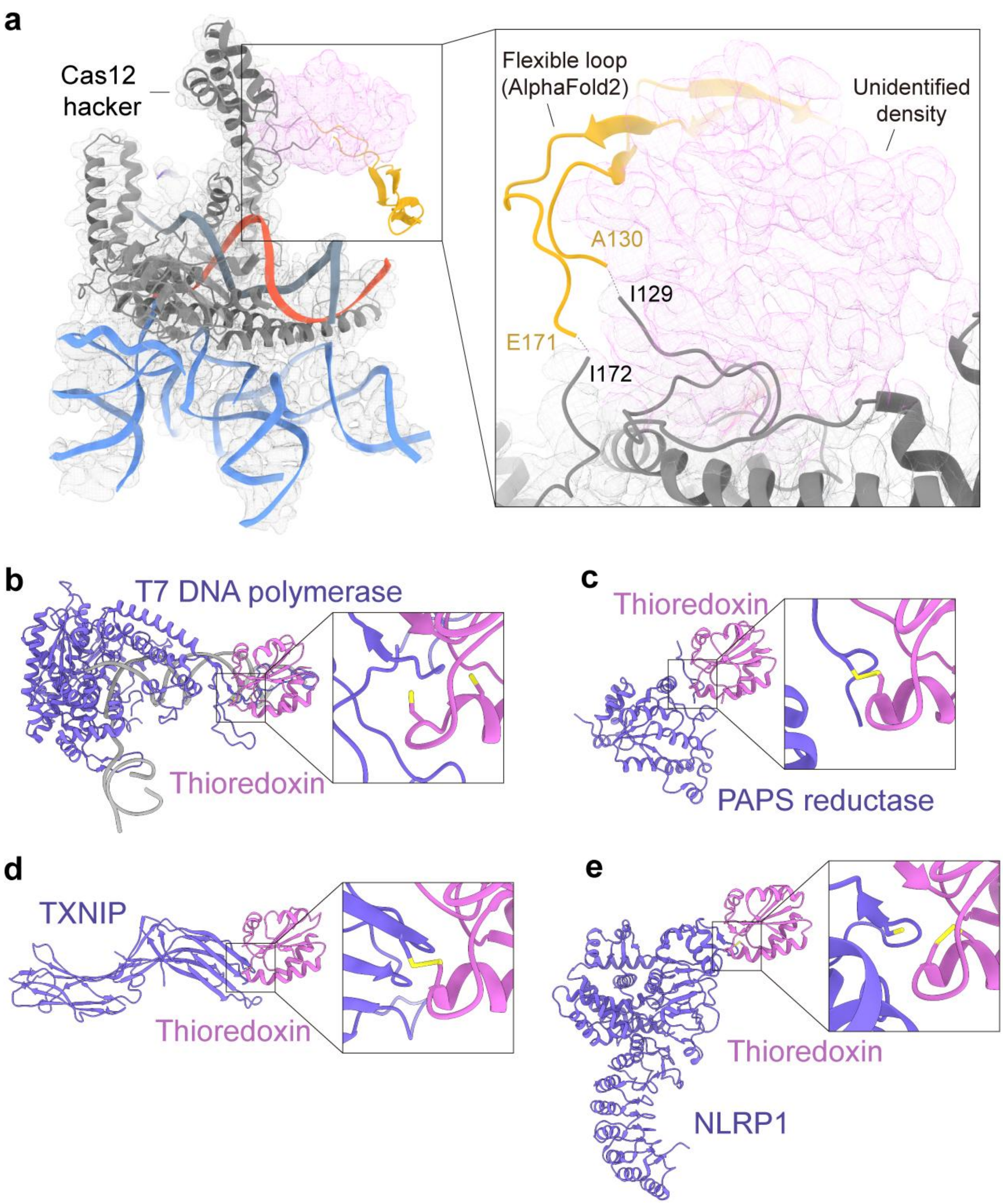


**Supplementary Fig. 5 Complex structures of different proteins and thioredoxin**

**a**, Close-up view of the predicted loop structure (goldenrod) and the unexpected globular density (violet mesh) of Cas12 hacker-sgRNA-target DNA (29-nt TS and 11-nt NTS) complex.

**b-e,** Structures of the T7 DNA polymerase-thioredoxin-primer/template DNA complex (PDB ID: 6N7W) (**b**), the PAPS reductase-thioredoxin complex (PDB ID: 2O8V) (**c**), the TXNIP-thioredoxin complex (PDB ID: 4LL1) (**d**), and the NLRP1-thioredoxin complex (PDB ID: 7WGE) (**e**). The thioredoxins are shown in orchid.

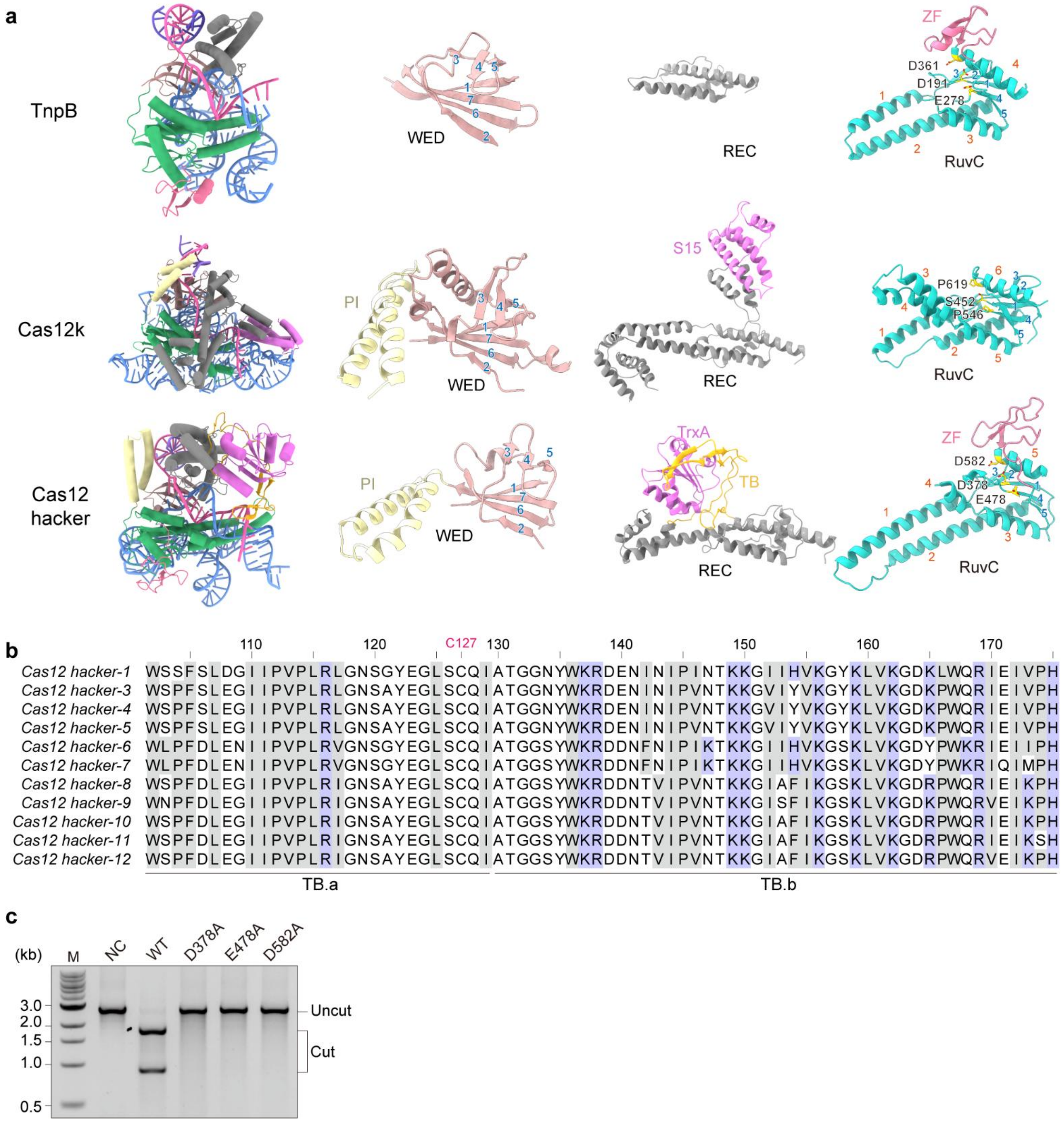


**Supplementary Fig. 6 Comparison of different domains in TnpB, Cas12k, and Cas12 hacker**

**a**, Structural comparison for WED, REC, and RuvC domains of TnpB (PDB ID: 8H1J), Cas12k (PDB ID: 8EA4), and Cas12 hacker. The WED, REC, and RuvC domains are represented in rosy brown, grey, and light sea green, respectively. α helices and β strands in the WED and RuvC domains are numbered in orange and blue, respectively.

**b**, Comparative sequence alignment of TB domains across the Cas12 hacker family, demonstrating the hydrophobic amino acids enrichment within TB.a and the prevalence of positively charged residues within TB.b. The hydrophobic and basic amino acids are shown in grey and purple background, respectively.

**c**, Probing the catalytic competence of three RuvC domain mutants (D378A, E478A, and D582A) within Cas12 hacker using the *in vitro* DNA cleavage assay. Each mutation individually results in undetected cleavage activity.

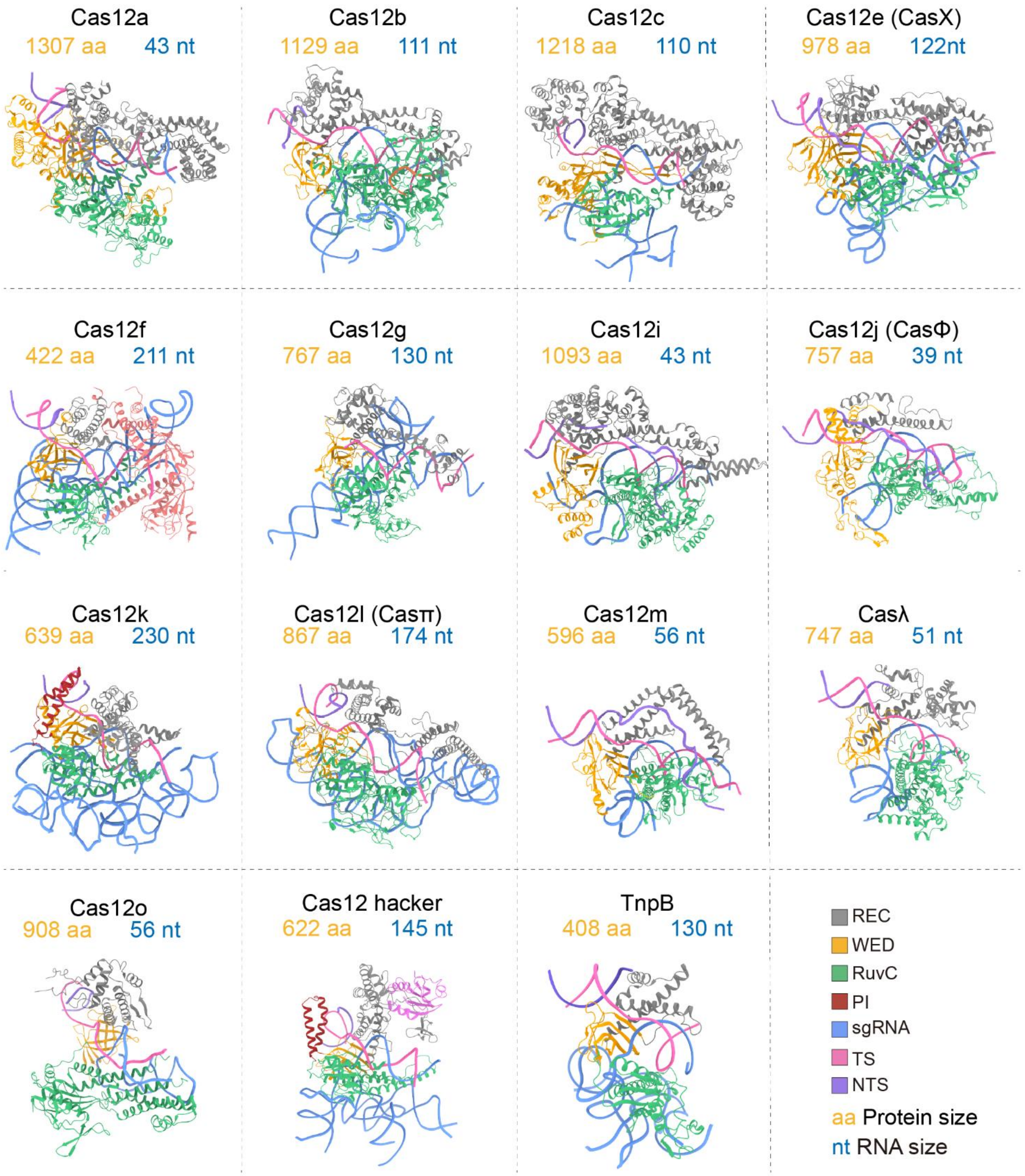


**Supplementary Fig. 7 Structural comparison of TnpB and Cas12 proteins in complex with corresponding sgRNA/ωRNA and target DNA**

Complex structures for Cas12a (PDB ID: 5B43), Cas12b (PDB ID: 5U31), Cas12c (PDB ID: 7V94), Cas12e (PDB ID: 7WAY), Cas12f (PDB ID: 8J12), Cas12g (PDB ID: 6XMG), Cas12i (PDB ID: 7D2L), Cas12j (PDB ID: 7LYT), Cas12k (PDB ID: 8EA4), Cas12l (PDB ID: 7YOJ), Cas12m (PDB ID: 8HHL), Casλ (PDB ID: 8DC2), Cas12o (PDB ID: 8XCA), TnpB (PDB ID: 8H1J), and Cas12 hacker.

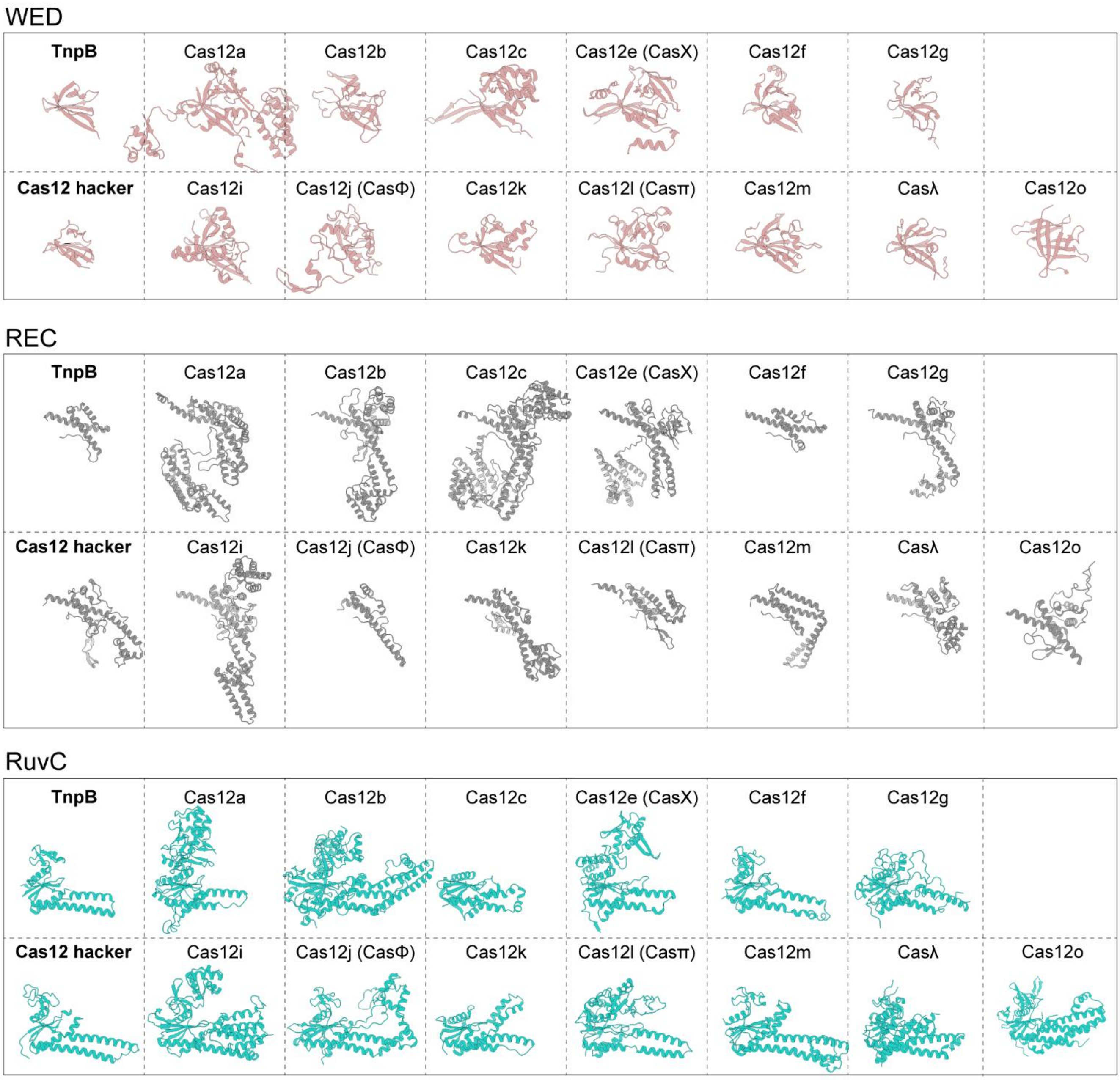


**Supplementary Fig. 8 Structural comparison of WED, REC, and RuvC domains for TnpB, Cas12 hacker, and other Cas12 proteins**

The TnpB (PDB ID: 8H1J), Cas12a (PDB ID: 5B43), Cas12b (PDB ID: 5U31), Cas12c (PDB ID: 7V94), Cas12e (PDB ID: 7WAY), Cas12f (PDB ID: 8J12), Cas12g (PDB ID: 6XMG), Cas12i (PDB ID: 7D2L), Cas12j (PDB ID: 7LYT), Cas12k (PDB ID: 8EA4), Cas12l (PDB ID: 7YOJ), Cas12m (PDB ID: 8HHL), Casλ (PDB ID: 8DC2), Cas12o (PDB ID: 8XCA), and Cas12 hacker proteins are roughly classified into WED (top, rosybrown), REC (middle, grey), and RuvC (bottom, lightseagreen) domains.

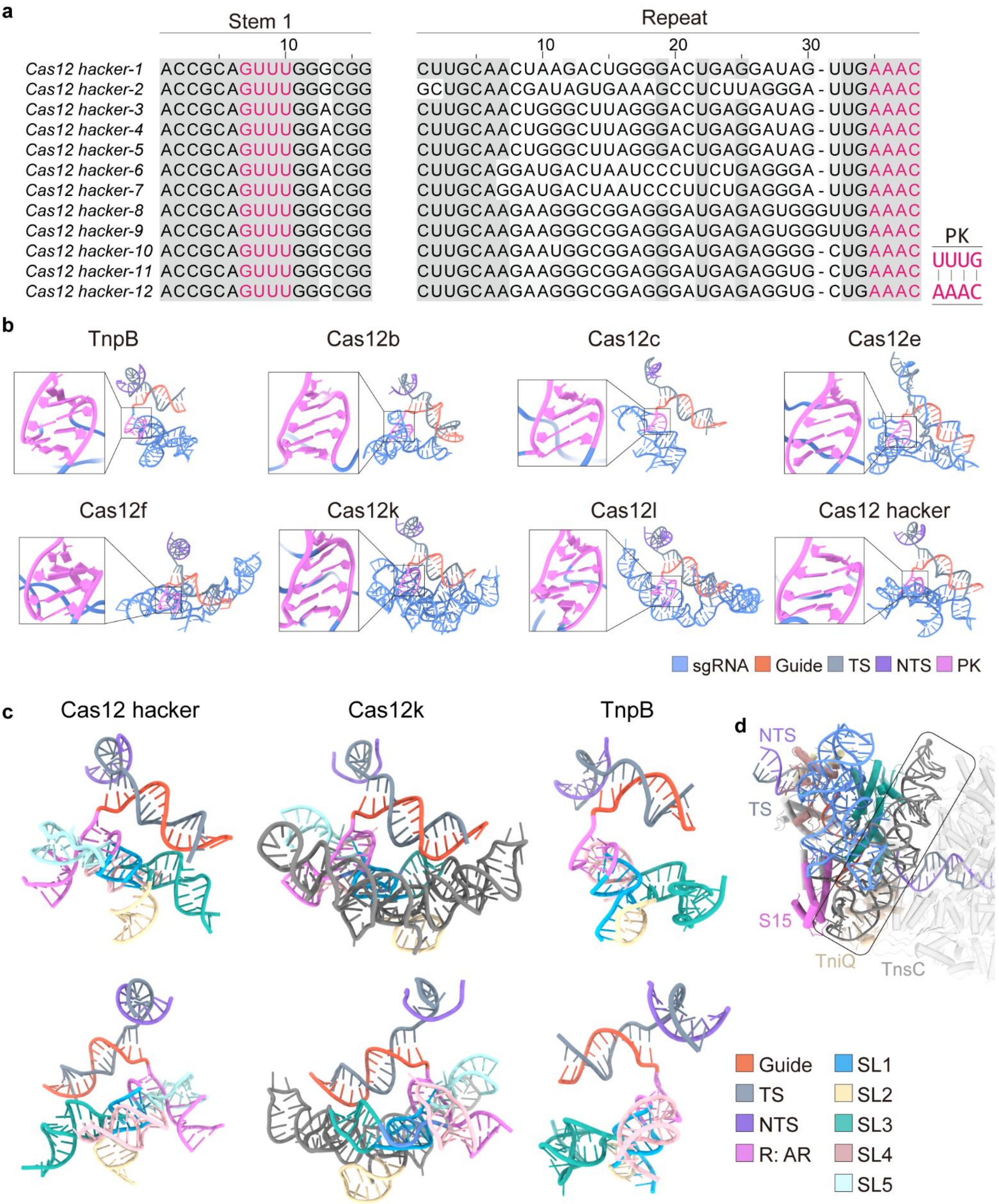


**Supplementary Fig. 9 Comparative analysis of sgRNA-target DNA architecture**

**a**, Sequence alignment comparing stem 1 (left) and the repeat sequence (right) across the Cas12 hacker family. The sgRNAs of this family exhibit a conserved AAAC:UUUG base-pairing pattern, essential for pseudoknot (PK) formation. The key AAAC:UUUG base-pairings are highlighted in violet.

**b**, Structural overview of the RNA-target DNA heteroduplex across various CRISPR-Cas12 systems, including TnpB (PDB ID: 8H1J), Cas12b (PDB ID: 5U31), Cas12c (PDB ID: 7V94), Cas12e (PDB ID: 7WAY), Cas12f (PDB ID: 8J12), Cas12k (PDB ID: 8EA4), Cas12l (PDB ID: 7YOJ), and Cas12 hacker. PK structures are distinctly marked in violet for clarity.

**c**, Structural comparison of the sgRNA/ωRNA-target DNA heteroduplexes of Cas12 hacker, Cas12k, and TnpB. The additional RNA elements in the sgRNA of Cas12k are indicated in dark grey.

**d**, Structure of the Cas12k-S15-TniQ-TnsC-sgRNA-target DNA complex. The extended sgRNA components (grey) that bridge Cas12k and Tn7 transposases are placed in the black box.

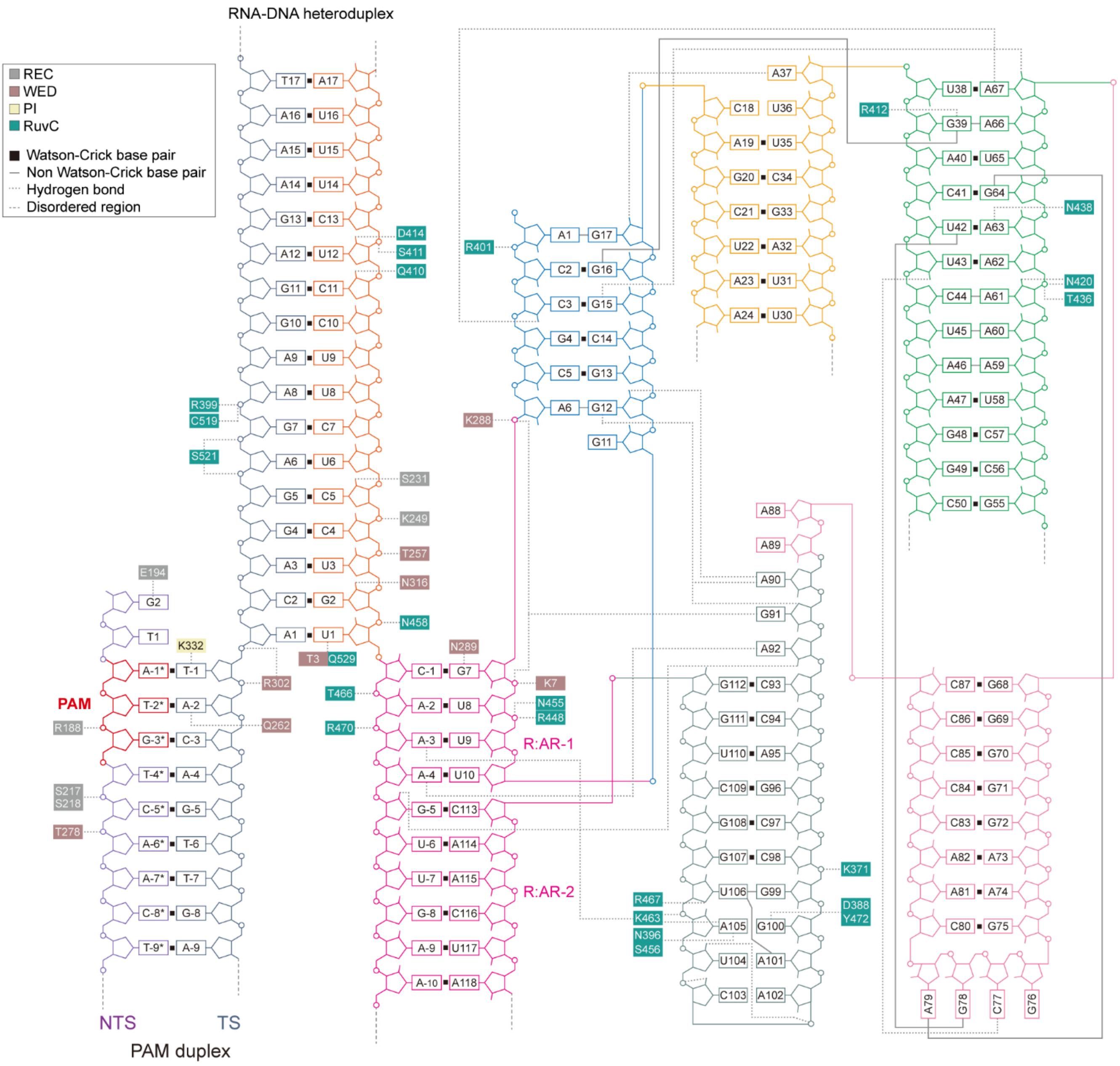


**Supplementary Fig. 10 Schematic for guide RNA and target DNA recognition**

The residues that interact with nucleic acids are placed in the coloured boxes.

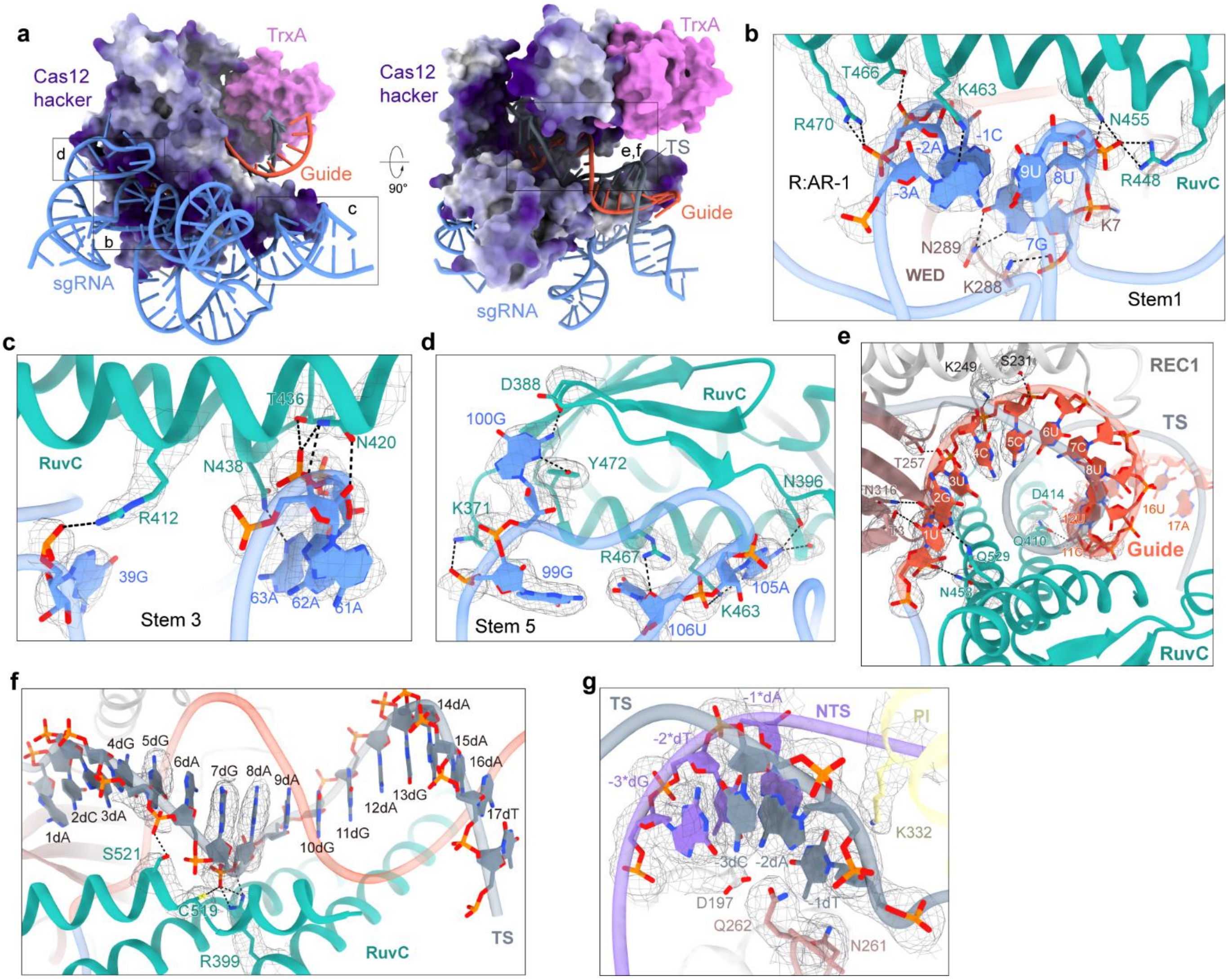


**Supplementary Fig. 11 Recognition of sgRNA and target DNA by Cas12 hacker**

**a**, Illustration of Cas12 hacker depicted via its electrostatic surface potential model, highlighting the interaction of sgRNA with the positively charged interface formed by the RuvC and WED domains. The sgRNA-target DNA heteroduplex is nestled within a positively charged central channel delineated by the REC and RuvC domains.

**b-d**, Interactions between Cas12 hacker's RuvC domain and sgRNA stem 1 (**b**), stem 3 (**c**), and stem5 (**d**). Hydrogen bonds are indicated by dashed lines.

**e,f**, Interactions of Cas12 hacker with the guide (**e**) and TS (**f**). Hydrogen bonds are indicated by dashed lines.

**g**, Cryo-EM density maps for the PAM duplex.

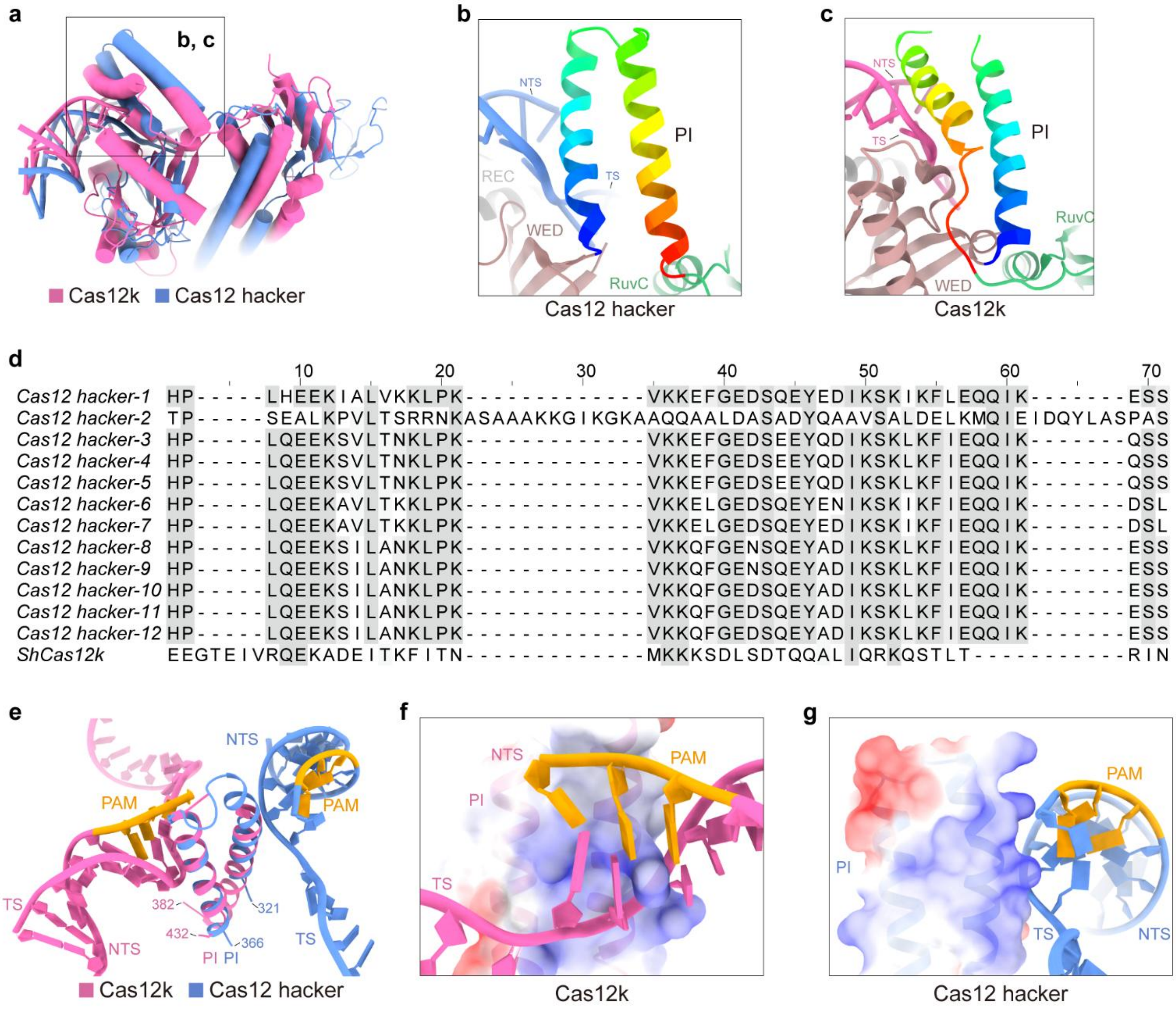


**Supplementary Fig. 12 Comparison of the PI domain of Cas12 hacker and Cas12k**

**a**, Structure superposition of the PI domains of Cas12k and Cas12 hacker.

**b,c,** Close-up view of the PI domain of Cas12 hacker (**b**) and Cas12k (**c**). The PI domains are visualized with a rainbow colour palette, transitioning from the blue N-terminus to the red C-terminus.

**d**, Multiple sequence alignment for PI domain of all Cas12 hacker nucleases and ShCas12k.

**e,** Structural alignment for PI domains of Cas12 hacker and ShCas12k with their corresponding target DNA.

**f,g**, Electrostatic surface potential of the PI domain of Cas12k (**f**) and Cas12 hacker (**g**) with corresponding target DNA.

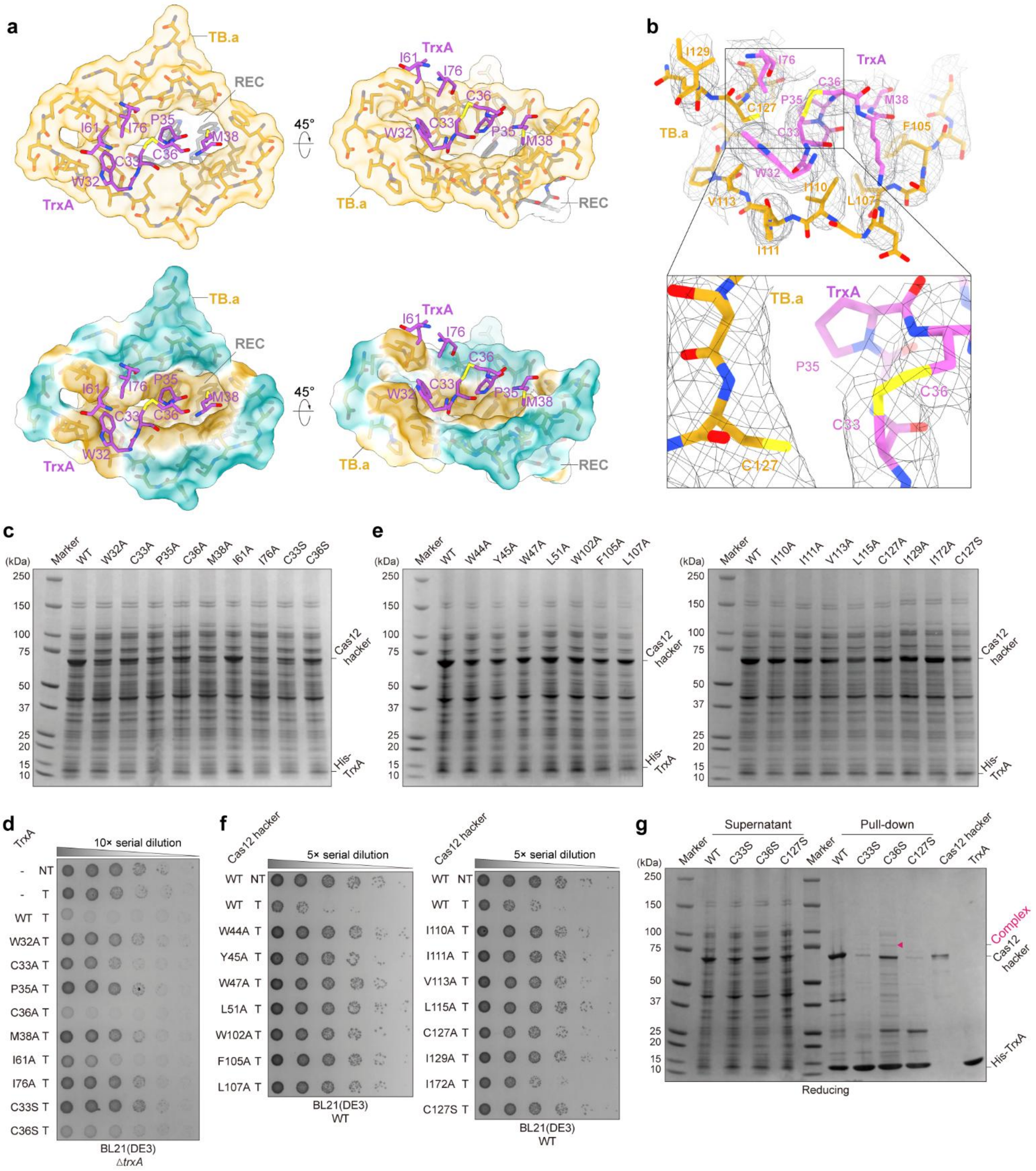


**Supplementary Fig. 13 Characterization of the interaction between TrxA and Cas12 hacker**

**a**, Illustration of the hydrophobic pocket formed by Cas12 hacker's TB and REC domains.

**b**, Cryo-EM density map of key residues in the interaction interface of TB and TrxA (top), and TrxA C33-C36 disulfide bond and Cas12 hacker C127 residue (bottom).

**c,** SDS-PAGE results for supernatants of wild-type (WT) Cas12 hacker purified with wild-type or mutant His-TrxA, visualized with CBB staining.

**d**, Evaluation of the functionality of wild-type and *trxA* mutants in an *E. coli trxA*-deletion strain utilizing bacterial spot assays, with non-targeting (NT) or targeting (T) sgRNA.

**e,** SDS-PAGE results for supernatants of wild-type or mutant Cas12 hacker proteins purified with wild-type His-TrxA, visualized with CBB staining.

**f**, Functional assessments of wild-type and mutant Cas12 hacker proteins in *E. coli* WT strain using bacterial spot assays with non-targeting (NT) or targeting (T) sgRNA.

**g,** Analysis of intermolecular disulfide bond between wild-type Cas12 hacker and wild-type or mutant forms of TrxA, as well as the Cas12 hacker (C127S) mutant with wild-type TrxA. The results were analyzed under reducing SDS-PAGE and stained with CBB. Data indicate that the intermolecular disulfide bond, observed in the TrxA (C36S) mutant and wild-type Cas12 hacker complex under non-reducing conditions (**Fig. 4f**), is conspicuously reduced upon application of reducing PAGE, suggesting the disulfide bond’s sensitivity to the reducing environment.

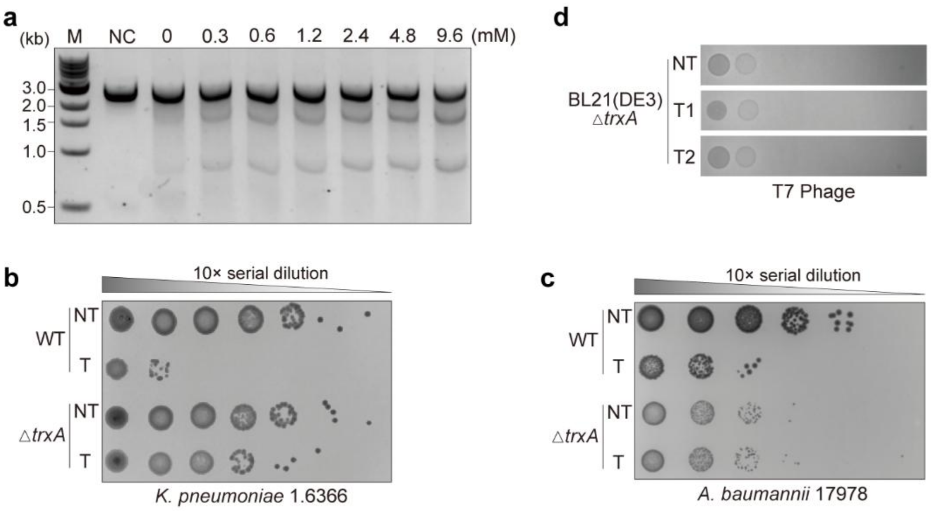


**Supplementary Fig. 14 The essential role of TrxA in modulating Cas12 hacker's activity**

**a**, *In vitro* cleavage assay for Cas12 hacker RNP complex purified from *trxA*-deletion *E. coli* BL21(DE3) with serially increasing the concentration of purified TrxA protein. The concentration of Cas12 hacker RNP is 300 nM.

**b**,**c**, Bacterial spot assays for Cas12 hacker targeting *Klebsiella pneumoniae* 1.6366 (**b**) and *Acinetobacter baumannii* 17978 (**c**) strains, with a non-targeting (NT) or targeting (T) sgRNA.

**d**, Phage plaque assays of T7 phage infecting the *E. coli* BL21 (DE3) *trxA*-deletion strain containing the CRISPR-Cas12 hacker system with a non-targeting (NT) or targeting (T) sgRNA.

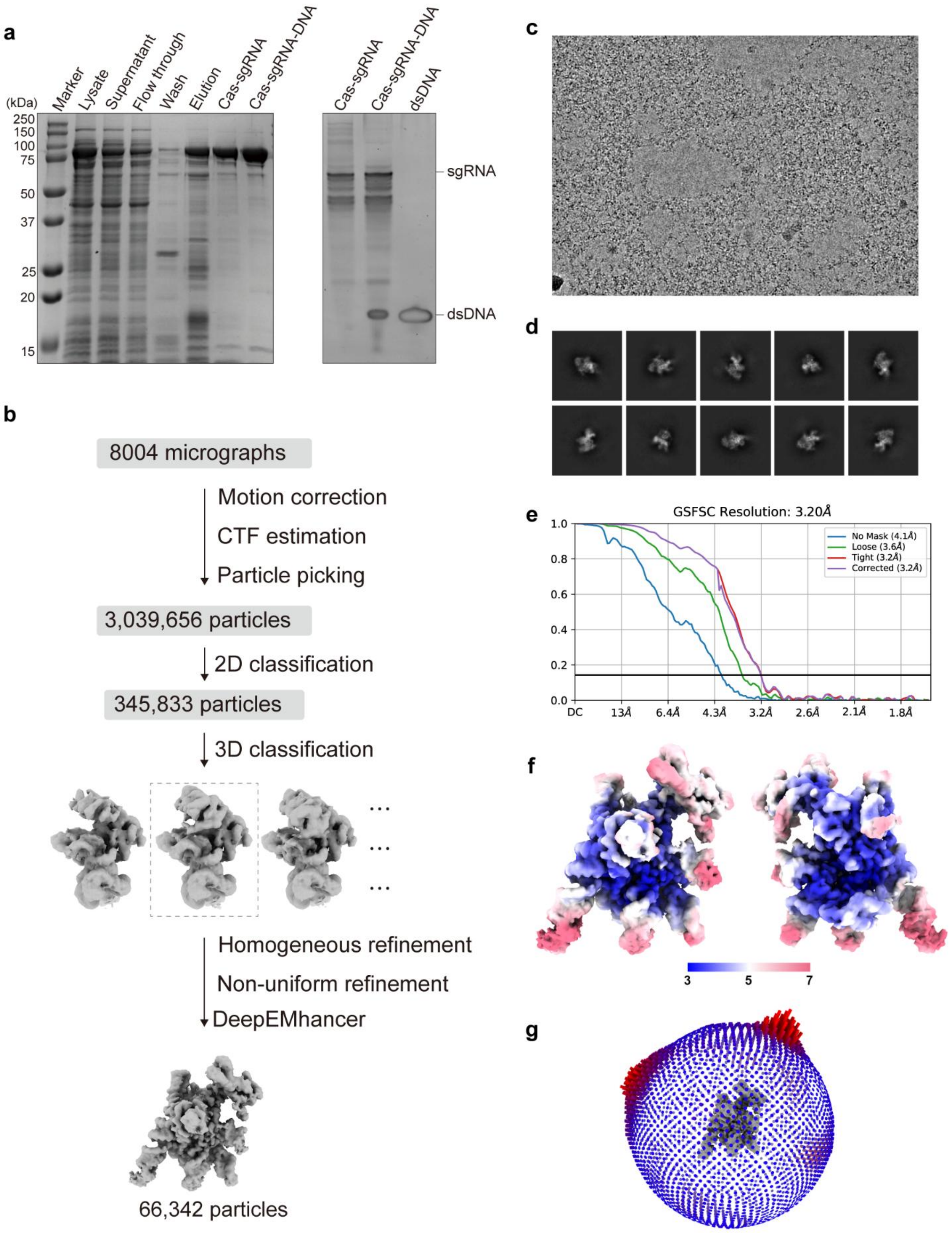


**Supplementary Fig. 15 Cryo-EM data analysis of the Cas12 hacker-TrxA-sgRNA-target DNA (33-bp dsDNA) complex**

**a**, SDS-PAGE (stained with CBB, left) and Urea-PAGE (right) analysis of the Cas12 hacker-sgRNA-target DNA (33-bp dsDNA) complex.

**b**, Cryo-EM data processing workflow for the Cas12 hacker-TrxA-sgRNA-target DNA (33-bp dsDNA) complex.
**c**, Representative cryo-EM micrograph from a total of 8004 micrographs.
**d**, Representative 2D class averages.
**e**, The half-map Fourier shell correlation (FSC) curves.
**f**, Euler angle distribution of particles.
**g**, Cryo-EM density map coloured by local resolution.

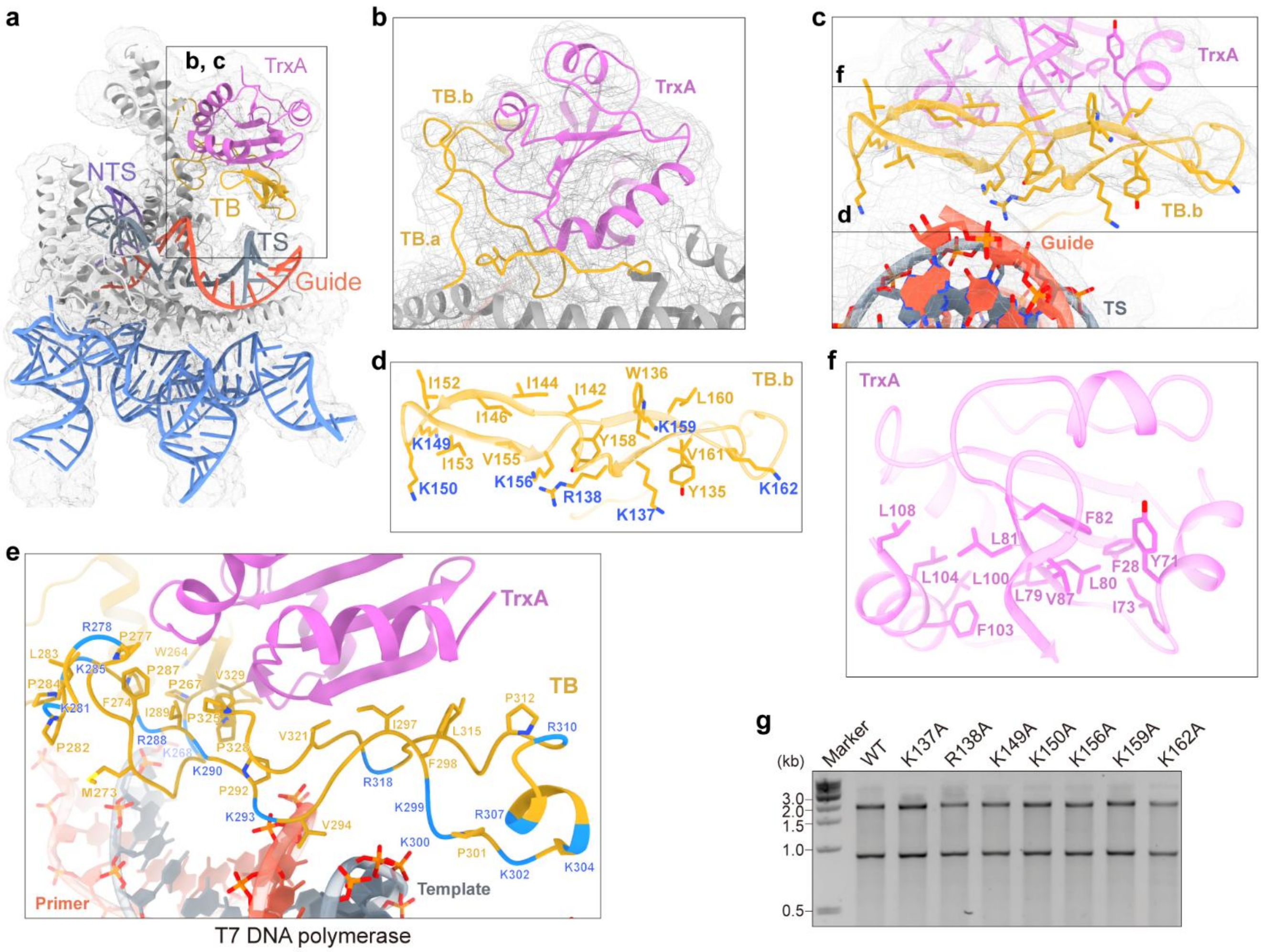


**Supplementary Fig. 16 The mechanism of Cas12 hacker's TB domain**

**a**, Cryo-EM density map and atomic model of the Cas12 hacker-TrxA-sgRNA-target DNA (33-bp dsDNA) complex.

**b**, Cryo-EM density map and atomic model of TrxA and the TB domain in the Cas12 hacker-TrxA-sgRNA-target DNA (33-bp dsDNA) complex.

**c**, Close-up view of interaction between TB.b and TrxA/guide-TS duplex in the Cas12 hacker-TrxA-sgRNA-target DNA (33-bp dsDNA) complex.

**d**, Close-up view of the TB.b domain of Cas12 hacker. Positively charged amino acids are shown in blue font and hydrophobic amino acids are shown in yellow font.

**e**, Close-up view of the TB domain of T7 DNA polymerase. Positively charged amino acids are shown in blue font and hydrophobic amino acids are shown in yellow font.

**f**, Close-up view of the interface of TrxA in interaction with TB.b. The interface contains many hydrophobic amino acids.

**g**, Comparative *in vitro* DNA cleavage assays for Cas12 hacker with wild-type (WT) and mutant TB.b domain. single mutation (K137A, R138A, K149A, K150A, K156A, K159A, or K162A) revealed unaltered cleavage activity compared with the wild-type Cas12 hacker.
